# Regulation of astrocyte metabolism by mitochondrial translocator protein 18kDa

**DOI:** 10.1101/2023.09.29.560159

**Authors:** Wyn Firth, Josephine L Robb, Daisy Stewart, Katherine R Pye, Rosemary Bamford, Asami Oguro-Ando, Craig Beall, Kate LJ Ellacott

**Author notes:** **Corresponding author:** Kate LJ Ellacott. **Author contributions: Wyn Firth:** conceptualization, data curation and analysis, funding acquisition, investigation, writing – original draft, writing – review and editing. **Josephine L Robb:** conceptualization, data curation and analysis, investigation, writing – review and editing. **Daisy Stewart:** investigation, data curation and analysis, writing – review and editing. **Katherine R Pye:** investigation, writing – review and editing. **Rosemary Bamford:** supervision, writing – review and editing. **Asami Oguro-Ando:** supervision, writing – review and editing. **Craig Beall:** conceptualization, funding acquisition, supervision, writing – review and editing. **Kate L J Ellacott:** conceptualization, data curation, funding acquisition, supervision, writing – original draft, writing – review and editing.

## Abstract

The mitochondrial translocator protein 18kDa (TSPO) has been linked to a variety of functions from steroidogenesis to regulation of cellular metabolism and is an attractive therapeutic target for chronic CNS inflammation. Studies in the periphery using Leydig cells and hepatocytes, as well as work in microglia, indicate that the function of TSPO may vary between cells depending on their specialised roles. Astrocytes are critical for providing trophic and metabolic support in the brain as part of their role in maintaining brain homeostasis. Recent work has highlighted that TSPO expression increases in astrocytes under inflamed conditions and may drive astrocyte reactivity. However, relatively little is known about the role TSPO plays in regulating astrocyte metabolism and whether this protein is involved in immunometabolic processes in these cells. Using TSPO-deficient (TSPO^-/-^) mouse primary astrocytes *in vitro* (MPAs) and a human astrocytoma cell line (U373 cells), we performed metabolic flux analyses. We found that loss of TSPO reduced basal astrocyte respiration and increased the bioenergetic response to glucose reintroduction following glucopenia, while increasing fatty acid oxidation (FAO). Lactate production was significantly reduced in TSPO^-/-^ astrocytes. Co-immunoprecipitation studies in U373 cells revealed that TSPO forms a complex with carnitine palmitoyltransferase 1a, which presents a mechanism wherein TSPO may regulate FAO in astrocytes. Compared to TSPO^+/+^ cells, inflammation induced by 3h lipopolysaccharide (LPS) stimulation of TSPO^-/-^ MPAs revealed attenuated tumour necrosis factor release, which was enhanced in TSPO^-/-^ MPAs at 24h LPS stimulation. Together these data suggest that while TSPO acts as a regulator of metabolic flexibility in astrocytes, loss of TSPO does not appear to modulate the metabolic response of astrocytes to inflammation, at least in response to the stimulus/time course used in this study.

## Introduction

The mitochondrial translocator protein 18kDa (TSPO) is a pentameric transmembrane protein with a high degree of evolutionary conservation^1–3^ that resides on the outer mitochondrial membrane of mammalian cells^4,5^, including glia^6^. TSPO has been linked to various functions including cholesterol transport into the mitochondria^3–5^, steroidogenesis^1,3–5^, inflammatory responses^5,7,8^, phagocytosis^9–11^, reactive oxygen species production^12,13^, and regulation of cellular metabolism^5,14,15^. This array of functions is highly relevant to glial cells, and while there is some knowledge about the role of TSPO in microglia, less is known about its role in astrocytes.

The brain is a metabolically expensive system with the highest weight-to-energy-use ratio of any organ in the body^16^. It is therefore essential that energy use in the CNS is tightly controlled. Astrocytes play key roles in nutrient uptake and distribution^17–19^, forming intimate relationships with neurons to provide trophic support and modulate neuronal activity directly; in turn, experimental modulation of these cells is sufficient to alter behavioural outputs^19–21^. In concert with microglia, astrocytes participate in the CNS inflammatory response by detecting and repairing damage, though this is not considered to be their primary function^22–24^. As inflammation requires changes in cellular metabolism, this affects energy use within the CNS^25–27^. We have previously shown that acute and chronic inflammation differentially alters the metabolic phenotype of primary mouse astrocytes in culture, modulating their expression of proteins involved with nutrient sensing and uptake, such as glucose transporter 1 (GLUT1)^28^. Previous work has demonstrated that chronic inflammation in the CNS is linked to upregulated TSPO expression^29^, largely attributed to glial cells^9,10,30^. In addition to regulating inflammatory responses, TSPO has been implicated in the regulation of cellular metabolism in the CNS and the periphery. For example, in 2016, Tu *et al.*^31^ provided the first evidence that TSPO may act as a regulator of fatty acid oxidation (FAO) in MA-10 Leydig cells, demonstrating that in these cells TSPO deficiency upregulates gene expression of carnitine palmitoyltransferase 1a (*Cpt1a)*, which encodes the rate-limiting enzyme of FAO. This study also showed that overexpression of TSPO downregulates *Cpt1a* expression, suggesting a bidirectional relationship^31^. Using murine microglia, a related study showed that in addition to metabolic impairments, microglial activation in response to proinflammatory stimulation is impaired by loss of TSPO^9^. More recently, Fairley *et al*.^10^ demonstrated that TSPO regulates glycolytic enzyme function and phagocytosis in microglia. However, in GBM1B stem-like cells and U87MG glioma cells, TSPO knockdown has been shown to increase glycolysis and reduce mitochondrial respiration^30^. The heterogeneity in these findings implies that TSPO may play cell-type and state-dependent roles in regulating cellular metabolism. Studies on the role of TSPO in astrocytes are necessary because of the developmental and functional distinctions between astrocytes and microglia^32–34^ including differential expression of metabolic enzymes, such as CPT1a which is expressed to a much greater extent in astrocytes^34,35^. Their unique functions require astrocytes to have fine control of energy sensing and cellular metabolism to rapidly respond to stimuli and restore the homeostatic balance of the brain. TSPO has received attention as a potential therapeutic target for neuroinflammatory conditions given its potential role at the interface of inflammatory responses and metabolism (‘immunometabolism’) in glia^10,36,37^. While it was recently shown that the astrocyte response to inflammatory stimuli includes regulation of TSPO expression^38^, the role TSPO plays in regulating astrocyte basal metabolism, inflammatory responses, and the metabolic response of astrocytes to inflammatory stimulation remains unclear.

Our work tested the hypothesis that loss of TSPO impairs astrocyte glucose metabolism and enhances FAO metabolism under basal conditions. We investigated a possible mechanistic basis for this using co-immunoprecipitations, elucidating a mechanism by which TSPO may regulate astrocyte metabolism. The impact of TSPO deficiency on the metabolic responses of astrocytes to an inflammatory stimulus was also explored.

## Methods

Unless otherwise stated, reagents were obtained from Fisher Scientific (UK).

### Animal use and husbandry

All animal studies were conducted in accordance with the UK Animals in Scientific Procedures Act 1986 (ASPA) and study plans were approved by the institutional Animal Welfare and Ethical Review Body at the University of Exeter. Mice were group housed on a 12:12 light-dark cycle at 22 ± 2°C with *ad libitum* access to standard laboratory rodent diet (LabDiet [EU] Rodent diet 5LF2; LabDiet, London, UK) and water.

As detailed in Morrissey *et al*. 2021^39^, mating pairs of TSPO heterozygote (^+/-^) (Tspo^tm1b^(EUCOMM)^Wtsi^) mice were used to produce the homozygote knockout (TSPO^-/-^) and wildtype (TSPO^+/+^) littermate mice used in these studies. Mice were crossed >7 generations onto a C57BL/6J background.

Mating pairs of C57BL/6J mice (Charles River, UK) were bred to produce mouse pups from which primary astrocytes were isolated. Data from this strain is presented in Figures 7 and 8.

Offspring of both sexes were used in these studies.

### Genotyping

TSPO genotypes (^+/+^, ^+/-^, ^-/-^) of breeder mice and mouse primary astrocytes (MPAs) were determined by PCR as reported previously in Morrissey *et al*. 2021^39^ (Supplementary Figure 1).

### Isolation of primary astrocytes and general cell culture

All cell cultures were maintained in 25mM glucose DMEM (Dulbecco’s Modified Eagle’s Medium; catalogue # D5671, Merck, UK) with 10% (v/v) FBS (Foetal Bovine Serum; catalogue # 10270106), 8mM L-glutamine (catalogue # 11500626) and 2% (v/v) penicillin-streptomycin (catalogue # 11528876). Cultures were maintained in humidified incubators at 37°C with 5% CO_2_.

U373 astrocytoma cells were purchased from the European Collection of Authenticated Cell Cultures (U-373 MG (Uppsala) (ECACC 08061901)^40^. U373 astrocytoma cells that were not subject to previous genetic modification were maintained up to p30 for use in these studies. TSPO^-/-^ and empty vector wildtype control U373 astrocytoma cells were p50-60. Though not ideal, this was unavoidable due to expanding single-cell colonies following genetic modification with the CRISPR-Cas9 system.

Mouse primary astrocytes (MPAs) were isolated from cortical tissue of neonates (postnatal days 1-5) as previously published^28^. All plastic and glassware used to culture MPAs was coated with poly-L-lysine (PLL; 4µg/mL; catalogue # P1274, Merck Life Sciences (UK)). TSPO^-/-^ MPAs were grown in T25 flasks until confluent before being stored in liquid nitrogen (10% v/v dimethyl sulfoxide (DMSO; catalogue # D2650, Merck Life Sciences (UK)) until TSPO genotypes had been confirmed. Once thawed, MPAs of the same genotype were pooled together into a T75 flask and grown until 100% confluent. MPAs from wildtype C57BL/6J neonates were grown in T75 flasks until confluent prior to storage in liquid nitrogen (10% DMSO). Immunocytochemistry was used to assess astrocyte purity via presence of glial fibrillary acidic protein (GFAP) immunoreactivity, and mean astrocyte purity was determined to be 98.67% (Supplementary Figure 2; n=10 coverslips, 2-5 images per coverslip over 3-4 individual collections).

The day before experiments, MPAs or U373 cells were seeded in DMEM (catalogue # 11966) with 7.5mM glucose (catalogue # G7021, Merck Life Sciences (UK)), 10% (v/v) FBS, 8mM L-glutamine and 2% (v/v) penicillin-streptomycin. On the day of experiments involving treatments, cells were cultured in serum-free DMEM (11966) supplemented with 2.5mM glucose for 2h before treatments were applied in the same media. When performing metabolic flux analyses or measuring metabolite production, XF DMEM (pH 7.4) (catalogue # 103575-100; Agilent, UK) with 2.5mM glucose, 2.5mM sodium pyruvate, and 2mM L-glutamine was used.

### Generation of TSPO-deficient U373 cells

Guide RNAs for TSPO were designed against Exons 2 and 3 of *Tspo* (Ensembl transcript: ENST00000396265.4 TSPO-203) using Benchling (Benchling software, 2019^41^) and cloned into either a pSpCas9(BB)-2A-GFP (PX458; Addgene plasmid #48138) plasmid or a pU6-(BbsI)_CBh-Cas9-T2A-mCherry plasmid (Addgene plasmid #64324) according to the method published by Ran *et al*. (2013)^42^. Plasmids validated by Sanger sequencing (Genewiz, Azenta Life Sciences Ltd, UK) were co-transfected into U373 cells using Lipofectamine LTX alongside an empty vector (EV) control prior to clonal selection. TSPO genotype was confirmed by PCR and Western blotting (Supplementary Figure 3). See Supplementary Methods for full gRNA details and primer sequences.

### Bacterial transformations and U373 transfections

Plasmids (Myc-tagged TSPO (‘TSPO-Myc’; catalogue # RC220107, OriGene) or a pCMV EV control (catalogue # PS100001, OriGene)) were grown in chemically competent DH5α *E. coli* following transformation via heat shock. 25µg/mL kanamycin (catalogue # K4000, Merck Life Sciences (UK)) was used to select successful transformants. Plasmids were extracted using a Plasmid Midi Kit (catalogue # 12143, Qiagen) according to the manufacturer’s specifications with the following modification: all centrifugation steps took place at 4600*xg* at room temperature and pressure (RTP) with appropriate modifications to the duration of spins (step 1: 15 minutes; step 10: 35 minutes; step 11: 15 minutes). Plasmids were allowed to air dry in a fume hood for 15 minutes before resuspension in 500µL Buffer TE (Qiagen). Plasmid concentration was quantified using a Nanodrop 2000.

U373 cells were transiently transfected in 6-well plates with 2.5µg plasmid (TSPO-Myc or pCMV EV) per well using Lipofectamine 3000 (catalogue # 15292465) according to the manufacturer’s specifications with minor modifications: transfection complexes were made up using serum-free D5671 DMEM with 8mM L-glutamine and 2% (v/v) penicillin-streptomycin 10 minutes before use. Cells in DMEM (catalogue # 11966) with 7.5mM glucose, 10% (v/v) FBS, 8mM L-glutamine and 2% (v/v) penicillin-streptomycin were seeded directly into transfection complex mixes and left for 24h before cell lysates were harvested.

### Cell treatments

Prior to treatment, cells were washed once with 0.01M phosphate-buffered saline (PBS; catalogue # 10209252) and incubated in serum-free 2.5mM glucose DMEM (catalogue # 11966) (‘assay media’) for 2h. Treatments were diluted to the appropriate concentrations in assay media immediately before application. 2-deoxyglucose (2DG; catalogue # D8375-5G, Merck Life Sciences UK; reconstituted in dH_2_O) was diluted to 10mM. Lipopolysaccharide from *Escherichia coli* (LPS; 026:B6; catalogue # L8274 Merck Life Sciences UK; reconstituted in dH_2_O) was diluted to a working concentration of 100ng/mL.

### Protein quantification

Cell lysates were collected using modified RIPA buffer (Supplementary Table 3) and manual scraping. Collected lysates were frozen, thawed on ice, and centrifuged at 21,000xg for 20 minutes at 4°C. Protein content of supernatant was quantified via Bradford assay according to the manufacturer’s instructions (catalogue # 500-0006, Bio-Rad). Absorbance was measured at 595nm using a Pherastar FS (BMG LABTECH).

### Co-immunoprecipitation

#### Immunoprecipitation of lysate from transfected cells

Co-immunoprecipitations were performed using Myc-Trap agarose beads (catalogue # yta-20, Proteintech) according to the manufacturer’s specifications with minor modifications: 20µL beads were used per tube; following the final wash/spin step, complexes were eluted in 2X SDS sample buffer (125mM Tris-HCl (pH 6.8), 4% SDS, 20% glycerol, bromophenol blue) by boiling at 95°C for 10 minutes. Eluate was centrifuged at 2500*xg* for 5 minutes at 4°C and samples were stored at −70°C until use in immunoblotting.

#### Immunoprecipitation of endogenous proteins from non-transfected cells

U373 cells were seeded at 3.0×10^6^ cells per dish in 150mm dishes. Cells were lysed with immunoprecipitation buffer (Supplementary Table 4) (4°C) and homogenised by turning 30 times in a Dounce homogeniser. Lysate was centrifuged at 1000*xg* for 10 minutes (4°C). 400µg protein was loaded per immunoprecipitation (IP). Protein G Dynabeads (catalogue # 10003D) were washed with 0.02% (v/v) PBS-Tween 20 (Tween-20: catalogue # P2285, Merck; 0.01M PBS) and incubated with 200µL of primary antibody mix (for antibody details see Supplementary Table 5) for 1h RTP on a rotary mixer. Supernatant was discarded and beads washed 3 times in 0.02% PBS-Tween prior to incubation with cell lysate diluted in 0.01% (v/v) n-dodecyl-β-maltoside (catalogue # 89902) in PBS (DBM-PBS; 0.01M PBS) on a rotary mixer for 2h at 4°C. Supernatant was moved to a clean tube and 50µL SDS sample buffer was added. This unbound fraction was incubated at 70°C for 5 minutes. Meanwhile, beads were washed three times with DBM-PBS, moved to a fresh tube, and bound protein was eluted using 50µL SDS sample buffer and heating at 70°C for 5 minutes. Immunoblotting was used to identify proteins of interest.

### Immunoblotting

Following sample quantification or co-immunoprecipitation, samples were run on 15% (v/v) polyacrylamide gels. In co-immunoprecipitation studies 20µL per fraction was loaded per well. For semi-quantifiable immunoblots, 10µg protein was loaded per well. Gels were run at 90V for 15 minutes, followed by 150V for 90 minutes. Gels were transferred onto nitrocellulose membranes via wet transfer at 100V for 70 minutes. Membranes were blocked using Odyssey blocking buffer (Tris-buffered saline [TBS]) (catalogue # 927-50000; Licor, UK) for 1h. Primary antibodies were applied overnight at 4°C or RTP for 1h. Membranes were washed 3 times in TBS (20mM Tris-HCl (pH 7.4), 152mM NaCl) with 0.05% (v/v) Tween-20; (TBS-T) before application of secondary antibodies. See Supplementary Methods (Supplementary Table 6) for antibody details. Membranes were scanned using an Odyssey CLx scanner (Licor, UK). Bands for proteins of interest were normalised to expression of GAPDH and data expressed as fold change over control.

### Immunocytochemistry

Cells were seeded onto PLL-coated coverslips at a density of 1×10^5^ cells per coverslip and left overnight. Cells were washed with 0.01M PBS and incubated (37°C, 5% CO_2_) in assay media for 2h prior to treatments or fixation. Cells were fixed by immersion in ice-cold 100% methanol for 90 seconds. Methanol was removed and cells were washed once with 0.01M PBS to prevent excessive dehydration. Coverslips were washed twice more with 0.01M PBS before blocking in 5% (v/v) normal donkey serum (NDS; catalogue # S30-100ml, Sigma Aldrich, UK) in 0.01M PBS (3% Tween-20) (PBS-T) for 15 minutes at RTP. Coverslips were then incubated with primary antibodies for 15 minutes (RTP) before being stored at 4°C overnight (18-20h). Primary antibodies were removed, and coverslips washed 3 times with PBS. Secondary antibodies were applied for 1h RTP in darkness. See Supplementary Methods (Supplementary Table 7) for antibody details. Coverslips were allowed to air dry in darkness before being mounted onto slides using Fluoroshield Mounting Medium with DAPI (catalogue # AB104139-20ML, Abcam, UK). Slides were imaged using a DM4000 B LED Fluorescent microscope.

### Metabolic flux analyses

Cells were seeded at a density of 4×10^4^ cells per well in 96 well plates 20-24h prior to experimentation. XF DMEM was used for all metabolic flux assays. Media was supplemented as appropriate for the assay used: for basal measurements of cellular metabolism in the presence of glucose and the mitochondrial stress test (MST), XF DMEM was supplemented with 2.5mM glucose (catalogue # G7021, Merck Life Sciences UK), 2.5mM sodium pyruvate (catalogue # P2256, Merck, UK), and 2mM L-glutamine (catalogue # 11500626); for the glycolysis stress test, XF DMEM was supplemented with 2mM L-glutamine; for measurements of cellular metabolism in the absence of glucose XF DMEM was supplemented with 2.5mM sodium pyruvate and 2mM L-glutamine. Prior to the assay, cells were washed once with XF DMEM (containing supplements relevant to the test used), which was immediately aspirated and replaced with more fresh media XF DMEM of an appropriate composition for the assay used. Cells were placed into a humidified non-CO_2_ incubator at 37°C to ‘degas’ for 60 minutes immediately prior to the assays. Following the degas period cells were placed into the XF^e^96 bioanalyzer and assays commenced. Readings were taken over 3-minute mix-measure cycles during the assay, with 3-4 measurements per cycle. A baseline read of 3-4 cycles was taken prior to any injections.

#### Mitochondrial stress test (MST)

The mitochondrial stress test was performed according to the manufacturer’s instructions (catalogue # 103015-100, Agilent, UK) with minor modifications. Following the initial baseline reads, oligomycin (0.5µM, Complex V inhibitor), carbonyl cyanide-4 (trifluoromethoxy) phenylhydrazone (FCCP; 1µM, OXPHOS uncoupler), and a rotenone-antimycin A mix (0.5µM, Complex I and III inhibitors) were injected sequentially to interrogate mitochondrial function. Parameters of the MST were calculated according to the manufacturer’s instructions.

#### Glycolysis stress test (GST)

This test was run in accordance with the manufacturer’s instructions (catalogue # 103020-100, Agilent, UK). Following the initial baseline reads, sequential injections of 10mM glucose (supraphysiological concentration to saturate the cells with glucose), 1µM oligomycin and 50mM 2-deoxyglucose (2DG; a non-metabolizable glucose analogue which competitively inhibits glycolysis by blocking the conversion of glucose to glucose-6-phosphate) were used to interrogate the glycolytic capabilities of the cells. Parameters of the GST were calculated according to the manufacturer’s instructions, substituting rates post-2DG for pre-glucose injection rates as a measure of non-glycolytic metabolism.

#### Fatty acid oxidation MST

Cells were plated 48h prior to the assay. 24h before the assay, plating media was replaced with fatty acid oxidation media (111mM NaCl, 4.7mM KCl, 1.25mM CaCl_2_, 2mM MgSO_4_, 1.2mM NaH_2_PO_4_, 2.5mM glucose, 0.5mM carnitine, and 5mM HEPES, pH 7.4) overnight. Carnitine supplementation was used to promote fatty acid oxidation (FAO). 45 minutes into the degas period, half the cells were treated with 40µM etomoxir, an inhibitor of CPT1 function. This concentration was chosen to avoid off-target effects associated with higher concentrations of etomoxir^43^. Immediately before running the assay, half of the etomoxir and control treated cells were treated with 200µM palmitate (0.17mM bovine serum albumin (BSA) vehicle catalogue # 10775835001, Merck, UK; 0.17mM BSA vehicle) or BSA vehicle (0.17mM, catalogue # 10775835001, Roche). An MST was then performed to assess mitochondrial fatty acid oxidation. Basal FAO was calculated as the difference in OCR between cells co-treated with palmitate and etomoxir, and cells treated with palmitate alone, immediately before oligomycin was injected. Maximal FAO was calculated as the difference in OCR between cells co-treated with palmitate and etomoxir, and cells treated with palmitate alone, at the highest rate measurement following FCCP injection.

#### Measurement of extracellular L-lactate

2.1×10^5^ cells were seeded in 6 well dishes 20-24h prior to experimentation. Following treatments, extracellular L-lactate levels were determined using the L-lactate Assay Kit (catalogue # 700510, Cayman Chemical, USA) according to the manufacturer’s specifications. Data are expressed as µM/mL.

#### Quantification of cytokine secretion

Following treatments, media samples were collected and centrifuged at 21,000xg for 5 mins (4°C). 1 volume of sample was diluted in 1 volume of assay media and stored at −70°C. Tumour necrosis factor (TNF) was quantified using a DuoSet ELISA kit (catalogue # DY410-05, R&D Systems, UK) according to the manufacturer’s instructions.

#### Cell viability assay

Cell viability was estimated using a propidium iodide stain (2µg/mL; catalogue # P4864, Merck Life Sciences (UK)) as previously published^28^.

### Data analysis

Microscope images were processed using FIJI (Fiji Is Just ImageJ^44^). Raw data were processed using Microsoft Excel and statistical analyses were performed using GraphPad Prism (v10.0.2). Extracellular flux analysis data were normalised to protein content per well using Wave Analysis Software (Agilent) and exported to Prism. Data are displayed as mean ± standard error of the mean. A result was deemed to be statistically significant if p<0.05.

## Results

### Loss of TSPO reduced astrocyte mitochondrial and non-mitochondrial respiration

To characterise the impact of loss of TSPO on astrocyte metabolism, TSPO^-/-^ MPAs and U373 cells were examined using the Seahorse XFe^96^ bioanalyzer (Figure 1). Based on published literature, we hypothesised that loss of TSPO would reduce astrocyte basal mitochondrial and glycolytic metabolism^9,31^. As anticipated, mean basal oxygen consumption rate (OCR; a proxy of mitochondrial metabolism) was reduced by 31.1% in TSPO^-/-^ MPAs compared to TSPO^+/+^ MPA controls (Figure 1A,B; p<0.0001). Similarly, mean basal extracellular acidification rate (ECAR; a proxy of glycolysis) was reduced by 38.8% (Figure 1C,D; p<0.0001). This led us to infer that under basal conditions TSPO^-/-^ MPAs showed reductions in aerobic glycolysis, and oxidative phosphorylation of its products, compared to TSPO^+/+^ counterparts. These results were replicated in TSPO^-/-^ U373 cells, where mean basal OCR was reduced by 15.7% (Figure 1E,F; p=0.0293) and mean basal ECAR was reduced by 45.6% compared to TSPO^+/+^ EV controls (Figure 1G,H; p<0.0001). Together, these data suggest that TSPO deficiency reduces the basal metabolic rates of astrocytes under basal conditions.

**Figure 1:**
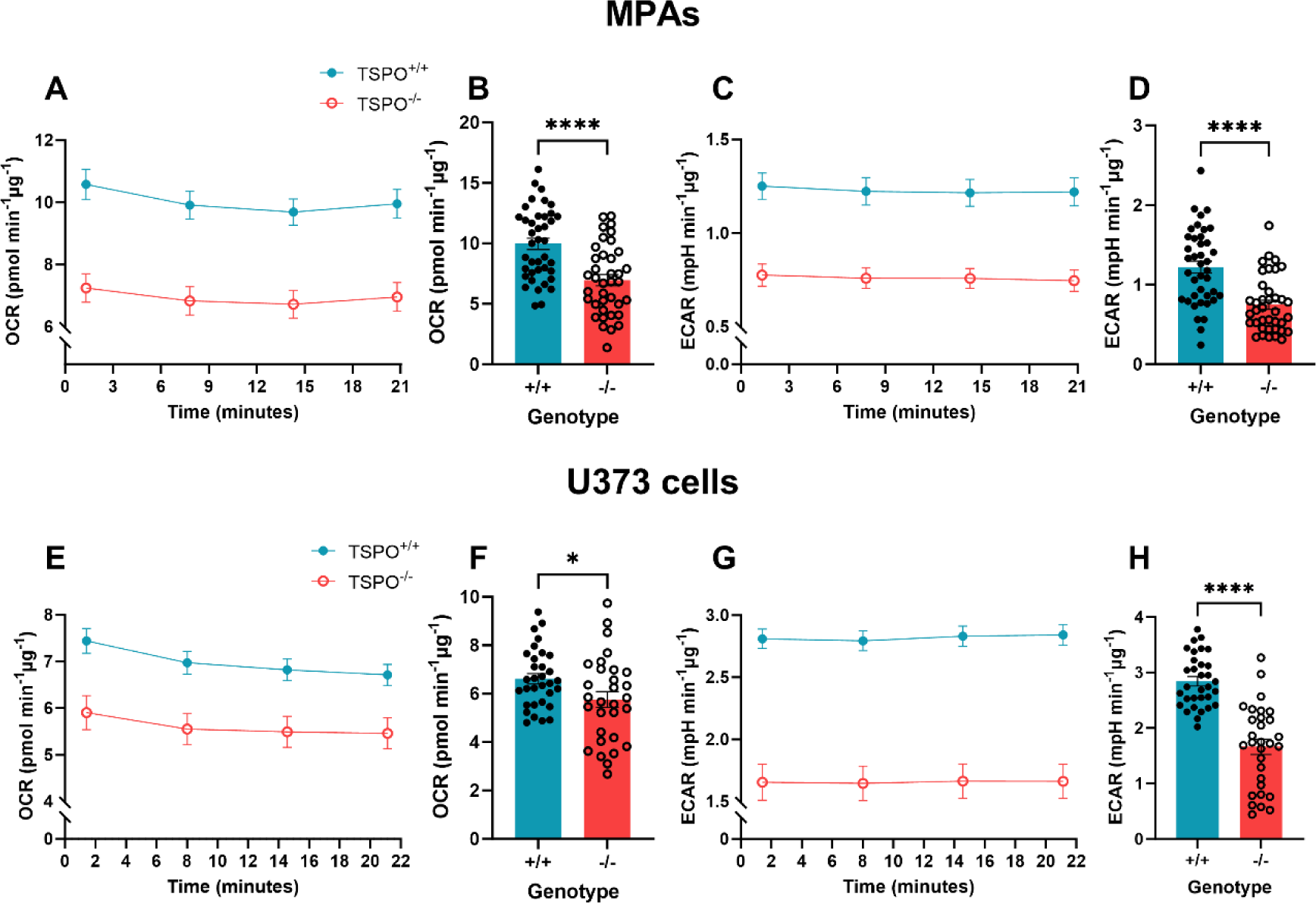
Loss of TSPO attenuated basal metabolism in astrocytes. **A:** Oxygen consumption rate (OCR) of TSPO-deficient (TSPO^-/-^) MPAs compared to wildtype (TSPO^+/+^) controls under basal conditions. **B:** Quantification of **A**, readings were taken from time point 4 of **A**. **C:** Extracellular acidification rate (ECAR) of TSPO^-/-^ MPAs under basal conditions. **D:** Quantification of **C**, readings were taken from time point 4 of **C**. n=38-40, data are pooled from across 3 independent plates. **E:** OCR of TSPO^-/-^ U373 cells compared to empty vector TSPO^+/+^ controls under basal conditions. **F:** Quantification of **E**, readings were taken from time point 4 of **E**. **G:** ECAR of TSPO^-/-^ U373 cells compared to empty vector TSPO^+/+^ controls under basal conditions. **H:** Quantification of **G**, readings were taken from time point 4 of **G**. n=30-32, data are pooled from across 3 independent plates. Unpaired two-tailed t-test. *p<0.05, ****p<0.0001. Data are expressed as mean ± standard error of the mean.

### TSPO deficiency attenuated the metabolic response to glucopenia

Once basal metabolic profiles of TSPO^-/-^ astrocyte models had been determined, we used the mitochondrial and glycolysis stress test paradigms to test the metabolic response to energetic stress in these cells, including the ability to switch to alternative substrates when glucose was limited.

We began by assessing changes to mitochondrial metabolism using the mitochondrial stress test (Figure 2A), and found that, in line with our initial data, basal mitochondrial respiration was significantly reduced in TSPO^-/-^ MPAs (Figure 2B, p=0.0006). This was accompanied by a reduction in mean maximal mitochondrial respiration of 40.2% compared to TSPO^+/+^ MPA controls (Figure 2C, p=0.0012). In TSPO^-/-^ MPAs mitochondria-linked ATP production was reduced by 35.8% (Figure 2D, p=0.0009), and mean proton leak was reduced by 46.0% (Figure 2E, p<0.0001). Coupling efficiency was increased by 2.3% (Figure 2F, p=0.0007), however mitochondrial spare capacity was reduced by 49.3% (Figure 2G, p=0.029). These results were mirrored in TSPO^-/-^ U373 cells (Figure 2H), where mean maximal respiration was reduced by 26.1% compared to TSPO^+/+^ EV controls (Figure 2I, p=0.0002 (basal); Figure 2J, p=0.001 (maximal)). Mitochondria-linked ATP production was reduced by 26.9% (Figure 2K, p=0.0004) and mean proton leak was reduced by 19.1% (Figure 2L, p=0.0011). In contrast to TSPO^-/-^ MPAs, there was no significant difference in mean coupling efficiency (Figure 2M, p=0.17) in TSPO^-/-^ U373 cells compared to TSPO^+/+^ EV controls, while – in keeping with TSPO^-/-^ MPAs – mean spare capacity was reduced by 27.3% (Figure 2N, p=0.0025). We found that under basal conditions U373 cells showed increased ECAR (a proxy of glycolysis) and reduced OCR (a proxy of mitochondrial respiration) relative to MPAs, therefore variation in the bioenergetic parameters between MPAs and U373 cells may be accounted for by these differences in basal bioenergetic profiles (Supplementary Figure 4).

**Figure 2:**
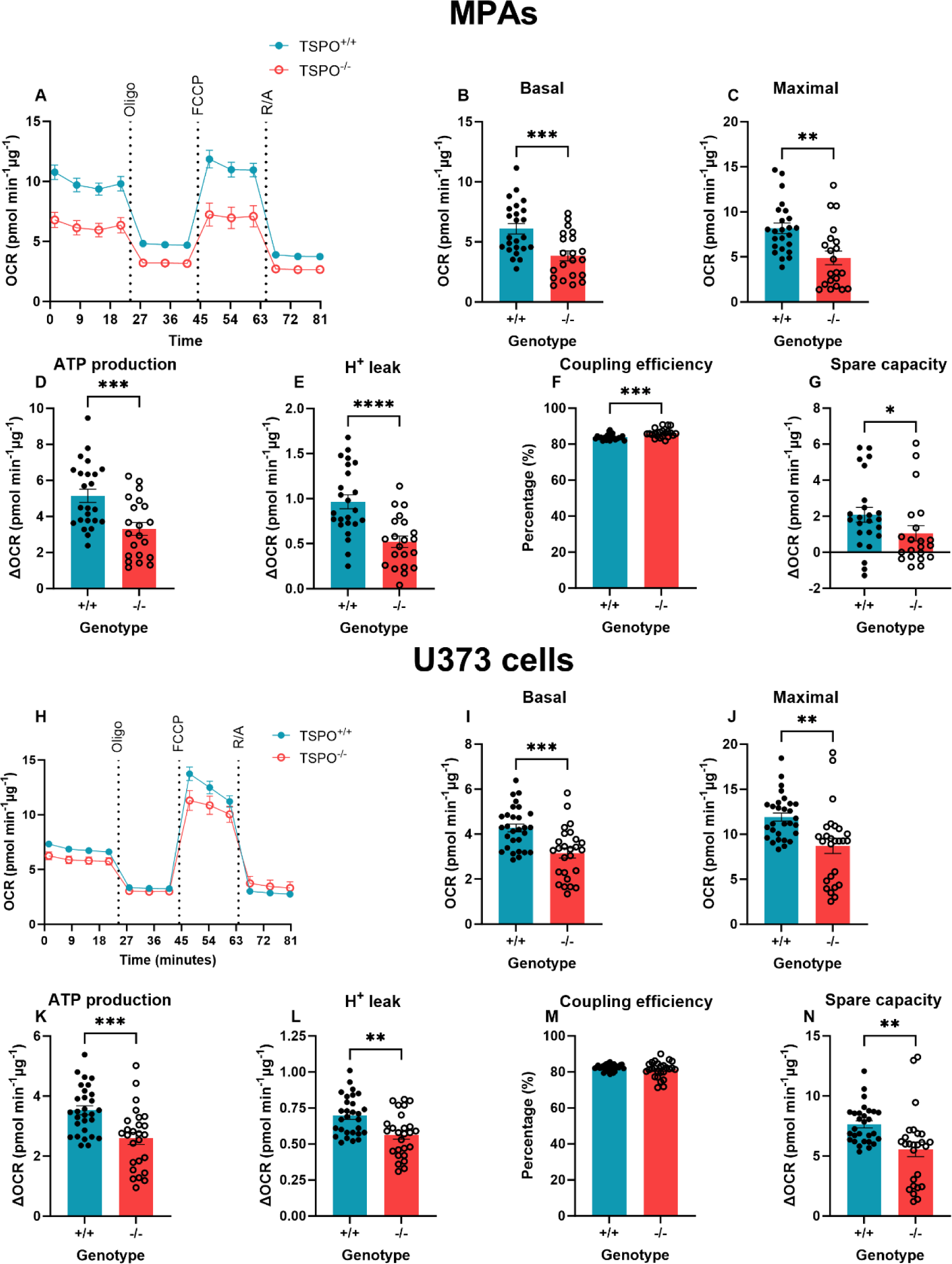
Mitochondrial respiration of astrocytes was reduced by TSPO deficiency. **A:** Oxygen consumption rate (OCR) of TSPO-deficient (TSPO^-/-^) MPAs compared to wildtype (TSPO^+/+^) controls during the mitochondrial stress test. Oligo = 0.5µM oligomycin, FCCP = 1µM carbonyl cyanide-p-trifluoromethoxyphenylhydrazone, R/A = 0.5µM rotenone/antimycin A. **B**: Basal mitochondrial respiration of TSPO^-/-^ MPAs compared to TSPO^+/+^ controls (difference in OCR prior to oligo injection and following R/A injection). **C:** Maximal mitochondrial respiration of TSPO^-/-^ MPAs compared to TSPO^+/+^ controls (difference in OCR following FCCP injection and after R/A injection). **D:** Mitochondria-linked ATP production of TSPO^-/-^ MPAs compared to TSPO^+/+^ controls (difference in OCR prior to and following oligo injection). **E:** Proton (H^+^) leak of TSPO^-/-^ MPAs compared to TSPO^+/+^ controls (difference in OCR after FCCP injection and R/A injection). **F:** Coupling efficiency of TSPO^-/-^ MPAs compared to TSPO^+/+^ controls (percentage of **B** used for ATP production). **G:** Mitochondrial spare capacity of TSPO^-/-^ MPAs compared to TSPO^+/+^ controls (difference in OCR between **B** and **C**). n=21-24 per genotype, data are pooled from across 3 separate plates. **H:** Oxygen consumption rate (OCR) of TSPO^-/-^ U373 cells compared to empty vector (EV) TSPO^+/+^ controls during the mitochondrial stress test. Oligo = 0.5µM oligomycin, FCCP = 1µM carbonyl cyanide-p-trifluoromethoxyphenylhydrazone, R/A = 0.5µM rotenone/antimycin A. **I**: Basal mitochondrial respiration of TSPO^-/-^ U373 cells compared to EV TSPO^+/+^ controls. **J:** Maximal mitochondrial respiration of TSPO^-/-^ U373 cells compared to EV TSPO^+/+^ controls. **K:** Mitochondria-linked ATP production of TSPO^-/-^ U373 cells compared to EV TSPO^+/+^ controls. **L:** Proton (H^+^) leak of TSPO^-/-^ U373 cells compared to EV TSPO^+/+^ controls. **M:** Coupling efficiency of TSPO^-/-^ U373 cells compared to EV TSPO^+/+^ controls (percentage of **I** used for ATP production). **N:** Mitochondrial spare capacity of TSPO^-/-^ U373 cells compared to EV TSPO^+/+^ controls (difference between **I** and **J**). n=26-29 per genotype, data are pooled from across 3 separate plates. Unpaired two-tailed t-test (**B-E, I-N**) or Mann-Whitney test (**F,G**). *p<0.05, **p<0.01, ***p<0.001, ****p<0.0001. Data are expressed as mean ± standard error of the mean.

Next, we examined the effect of TSPO deficiency on astrocyte glycolysis via the glycolysis stress test (GST; Figure 3A (MPAs), 3F (U373 astrocytoma cells)). In this paradigm, a glucose-free incubation period (glucopenia) followed by glucose injection is used to examine the glycolytic rate of the cells. Because our previous data (Figures 1 and 2) showed that TSPO deficient astrocytes had reduced ECAR, we hypothesised that these cells would exhibit reduced glycolysis. Non-glycolytic acidification – acidification of the media that may be attributed to other cellular or metabolic processes that produce H^+^, such as FAO^45^ – was significantly increased by 24.4% in TSPO^-/-^ MPAs (Figure 3B, p<0.0001). Meanwhile, in contrast to our expectations, we found that mean glycolytic rate was increased by 18.0% in TSPO^-/-^ MPAs (Figure 3C, p<0.0001). Similarly, mean glycolytic capacity and reserve were increased by 25.8% and 27.6% respectively (Figure 3D,E; p<0.0001 (both parameters)). In TSPO^-/-^ U373 cells (Figure 3F), non-glycolytic acidification was increased by 21.1% (Figure 3G, p=0.0057). In contrast to TSPO^-/-^ MPAs, in TSPO^-/-^ U373 cells, mean glycolytic rate was reduced by 26.5% (Figure 3H, p<0.0001) and we observed no change in glycolytic capacity in TSPO^-/-^ U373 cells compared to TSPO^+/+^ EV controls (Figure 3I, p=0.38). However, mean glycolytic reserve was increased by 72.9% in TSPO^-/-^ U373 cells (Figure 3J, p<0.0001).

**Figure 3:**
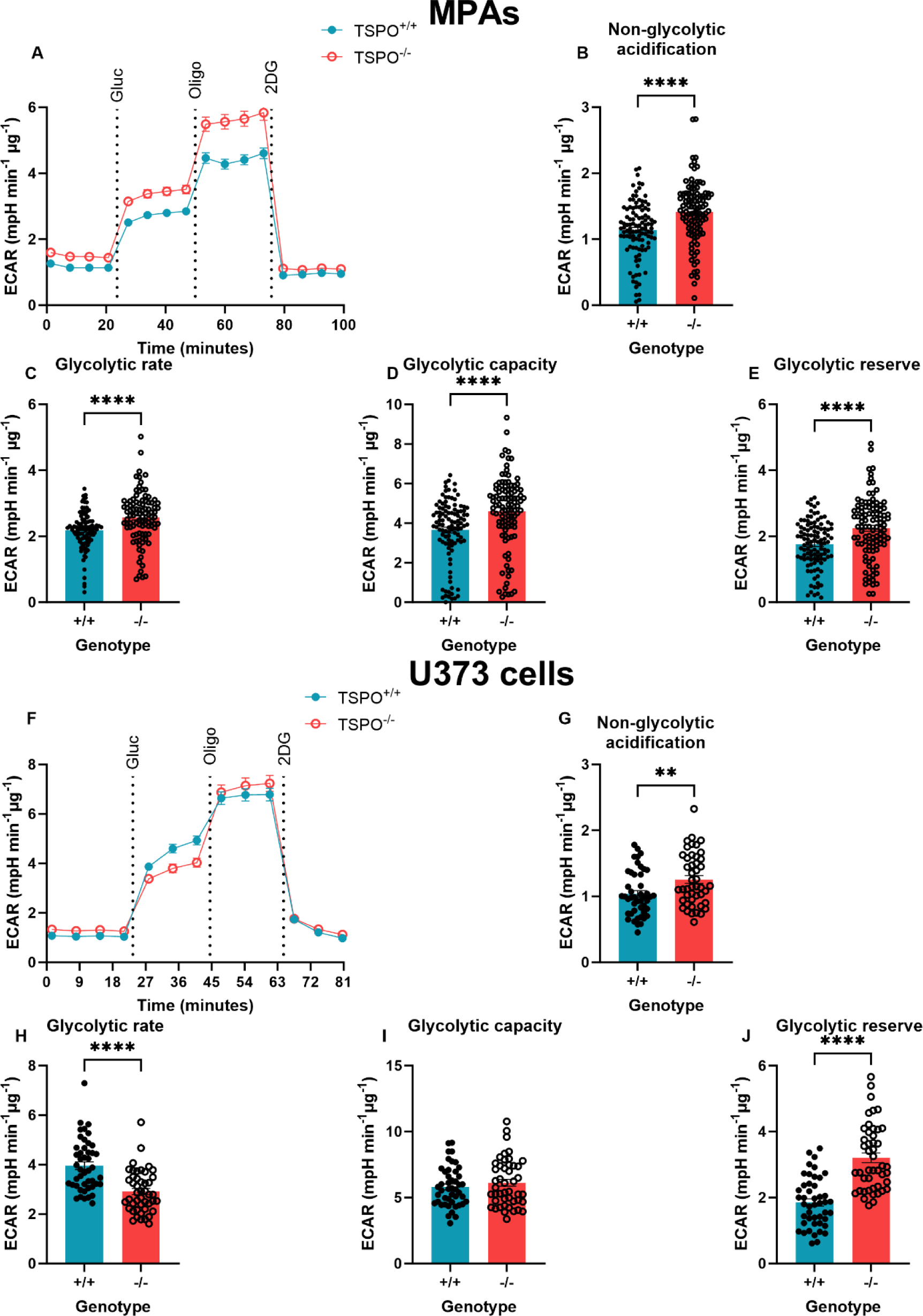
TSPO deficiency increased glycolytic rate in MPAs but reduced glycolytic rate in U373 astrocytoma cells. **A:** Extracellular acidification rate (ECAR) of TSPO-deficient (TSPO^-/-^) MPAs compared to wildtype (TSPO^+/+^) controls during the glycolysis stress test. Gluc = 10mM glucose, oligo = 1µM oligomycin, 2DG = 50mM 2-deoxyglucose. **B:** Non-glycolytic acidification of TSPO^-/-^ MPAs compared to TSPO^+/+^ controls (ECAR immediately prior to glucose injection). **C:** Glycolytic rate of TSPO^-/-^ MPAs compared to TSPO^+/+^ controls (ECAR after glucose injection immediately prior to oligo injection). **D:** Glycolytic capacity of TSPO^-/-^ MPAs compared to TSPO^+/+^ controls (difference between maximum ECAR after oligo injection and non-glycolytic acidification). **E:** Glycolytic reserve of TSPO^-/-^ MPAs compared to TSPO^+/+^ controls (difference between glycolytic rate and capacity). n=105-108, data are pooled from across 3 independent plates. **F:** ECAR of TSPO^-/-^ U373 cells compared to empty vector (EV) TSPO^+/+^ controls during the glycolysis stress test. Gluc = 10mM glucose, oligo = 1µM oligomycin, 2DG = 50mM 2-deoxyglucose. **G:** Non-glycolytic acidification of TSPO^-/-^ U373 cells compared to EV TSPO^+/+^ controls. **H:** Glycolytic rate of TSPO^-/-^ U373 cells compared to EV TSPO^+/+^ controls. **I:** Glycolytic capacity of TSPO^-/-^ U373 cells compared to EV TSPO^+/+^ controls. **J:** Glycolytic reserve of TSPO^-/-^ U373 cells compared to EV TSPO^+/+^ controls. n=46, data are pooled from across 3 independent plates. Unpaired two-tailed t-test (**B,E,G,I,J**) Mann-Whitney test (**C,D,H**). **p<0.01, ****p<0.0001. Data are expressed as mean ± standard error of the mean.

We were particularly intrigued by the apparently contradictory nature of our results in TSPO^-/-^ MPAs – whereas our initial data showed that a proxy of glycolysis (ECAR) was reduced in TSPO^-/-^ MPAs in the presence of glucose (Figures 1 and 2), our data from the glycolysis stress test (Figure 3) showed a relative increase in glycolytic metabolism in these cells, including basal ECAR prior to the injection of glucose (non-glycolytic acidification, Figure 3B). Postulating that this may have arisen due to a difference in the experimental protocol used – namely, the glucose starvation period followed by the reintroduction of glucose inherent to the GST paradigm – we hypothesised that TSPO deficiency may alter the metabolic adaptations when glucose is absent (glucopenia). We tested this by recapitulating the initial conditions of the MST (glucose present throughout) and GST (glucose absent initially) on the same plate: incubating TSPO^-/-^ and TSPO^+/+^ MPAs either in the presence or absence of glucose to directly compare their basal metabolic rates under these different conditions (Figure 4A-D). Using a two-way ANOVA we observed no statistically significant effects of genotype on OCR (Figure 4A,C; p_genotype_=0.64, F_(1,58)_ = 0.2246) and ECAR (Figure 4B,D; p_genotype_=0.16, F_(1,58)_ = 1.987). However, we observed a statistically significant impact of glucose concentration on OCR (Figure 4A,C; p_glucose_=0.044, F_(1,58)_ = 4.232) and ECAR (Figure 4A,D; p_glucose_=0.035, F_(1,58)_ = 4.654), and a statistically significant interaction of these variables on OCR (p_interaction_=0.0034, F_(1,58)_ = 9.348) and ECAR (p_interaction_=0.019, F_(1,58)_ = 5.186).

**Figure 4:**
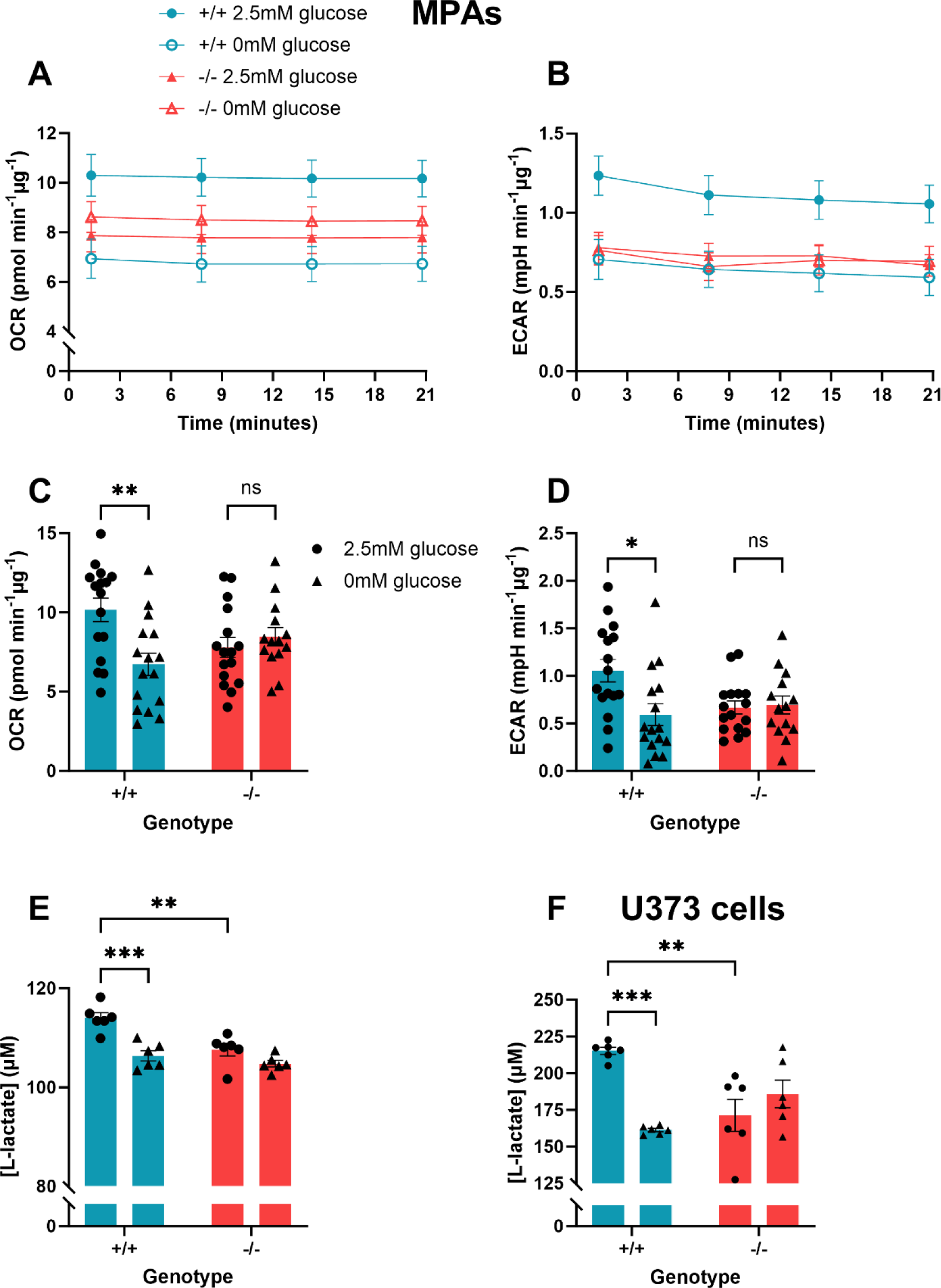
TSPO^-/-^ astrocytes secreted less L-lactate in the presence of glucose and lactate secretion was not affected by glucopenia. **A:** Oxygen consumption rate (OCR) of TSPO-deficient (TSPO^-/-^) MPAs ± glucose (2.5mM) compared to wildtype (TSPO^+/+^) controls ± glucose. **B:** Extracellular acidification rate (ECAR) of TSPO^-/-^ MPAs ± glucose compared to TSPO^+/+^ controls ± glucose. **C:** Quantification of **A** at time point 4. **D:** Quantification of **B** at time point 4. **E:** Extracellular L-lactate concentrations sampled from TSPO^-/-^ MPAs ± glucose and TSPO^+/+^ controls ± glucose. **F:** Extracellular L-lactate concentrations sampled from TSPO^-/-^ U373 cells ± glucose and empty vector TSPO^+/+^ controls ± glucose. **A-D:** n=14-16 from one plate to directly examine trends in basal metabolism observed in Figures 1-3. **C,D,E,F:** 2-way ANOVA with Šídák’s multiple comparisons test. **C:** p_genotype_=0.6373, F_(1,58)_ = 0.2246. p_glucose_=0.0442, F_(1,58)_ = 4.232. p_interaction_=0.0034, F_(1,58)_ = 9.348. **D:** p_genotype_=0.1640, F_(1,58)_ = 1.987. p_glucose_=0.0351, F_(1,58)_ = 4.654. p_interaction_=0.0191, F_(1,58)_ = 5.186. **E:** p_genotype_=0.0011, F_(1,20)_ = 14.55. p_glucose_ <0.0001, F_(1,20)_ = 25.04. p_interaction_=0.0322, F_(1,20)_=5.298. **F:** p_genotype_=0.2011, F_(1,20)_ = 1.748. p_glucose_=0.00145, F_(1,20)_ = 7.159. p_interaction_=0.0001, F_(1,20)_ = 21.84. n=6. ns p>0.05, *p<0.05, **p<0.01, ***p<0.001. Data are expressed as mean ± standard error of the mean.

Post-hoc analyses revealed that, under glucose-free conditions, mean basal OCR and ECAR of TSPO^+/+^ MPAs were reduced by 33.9% and 43.9% respectively (Figure 4A-D; p=0.0031 [OCR], p=0.0103 [ECAR]). Although the basal OCR and ECAR of TSPO^-/-^ MPAs were lower than TSPO^+/+^ MPAs they did not change in the absence of glucose (Figure 4A-D; p=0.9823 [OCR], p>0.9999 [ECAR]) suggesting that TSPO^-/-^ MPAs may be less dependent on glucose to maintain their basal metabolic rate than their TSPO^+/+^ controls.

To confirm the results of our extracellular flux analyses (Figure 4 A-D), we quantified the secretion of L-lactate from TSPO^-/-^ and wildtype control astrocytes as a secondary experimental measure of glycolysis (Figure 4E,F). As the conjugate base of lactic acid, L-lactate would acidify media if secreted from the cells, resulting in the changes in ECAR observed in our metabolic flux assays. In MPAs, TSPO genotype had a statistically significant effect on L-lactate secretion (Figure 4E p_genotype_=0.0011, F_(1,20)_ = 14.55), as did glucose concentration (p_genotype_<0.0001, F_(1,20)_ = 25.04). We observed a statistically significant interaction of these variables in MPAs (p_genotype_=0.032, F_(1,20)_ = 5.298). Post-hoc analysis revealed that in the presence of glucose TSPO^-/-^ MPAs secreted 5.6% less L-lactate than TSPO^+/+^ controls (Figure 4E, p=0.002). In the absence of glucose, while TSPO^+/+^ MPAs reduced their L-lactate secretion by 6.7% (Figure 4E, p=0.0003), L-lactate secretion was not altered in TSPO^-/-^ MPAs (Figure 4E, p=0.35). In U373 astrocytoma cells, TSPO genotype did not have a statistically significant effect on L-lactate secretion (Figure 4F, p_genotype_=0.20, F_(1,20)_ = 1.748), however we observed a statistically significant effect of glucose concentration on L-lactate secretion from these cells (p_glucose_=0.0015, F_(1,20)_ = 7.159). Moreover, we observed a statistically significant interaction between these variables (p_interaction_=0.0001, F_(1,20)_ = 21.84). Post-hoc analyses supported our results from MPAs: mean L-lactate secretion was reduced by 20.4% in TSPO^-/-^ U373 cells compared to TSPO^+/+^ controls (Figure 4F, p=0.0024) under basal conditions. Further, unlike TSPO^+/+^ controls, which altered their L-lactate secretion in response to glucopenia (Figure 4F, p=0.0003), L-lactate secretion from TSPO^-/-^ U373 cells was unchanged under glucopenic conditions (Figure 4F, p=0.68).

Together, our data from MPAs and U373 astrocytoma cells supports the notion that TSPO^-/-^ astrocytes are less reliant on glucose than TSPO^+/+^ controls to maintain their metabolic requirements in the absence of glucose and recapitulates the data we obtained using the metabolic flux analyses (Figure 4A-D). Thus, when considered with our metabolic flux analyses (Figure 4A-D), these data suggest that, compared to TSPO^+/+^ controls, TSPO^-/-^ astrocytes may be meeting their bioenergetic requirements by preferentially metabolising substrates other than glucose.

### TSPO^-/-^ astrocytes used fatty acid oxidation to maintain their metabolic rate

As we observed that TSPO^-/-^ astrocytes were able to maintain their metabolic parameters (albeit at a lower bioenergetic rate) in the absence of glucose (Figure 4A-D), and produced less L-lactate under basal conditions (Figure 4E,F), we postulated that this may be due to TSPO^-/-^ astrocytes being more readily able to utilise other metabolic substrates than TSPO^+/+^ controls.

In MA-10 Leydig cells, which like astrocytes are capable of fatty acid metabolism and steroidogenesis, loss of TSPO can increase fatty acid oxidation and overexpression of TSPO is linked to reduced expression of genes involved with FAO^31^. Hence, we hypothesised that TSPO^-/-^ astrocytes were meeting their bioenergetic requirements through enhanced fatty acid metabolism. We examined this using a fatty acid oxidation stress test (FAOST; Figure 5A,D). This assay involves an overnight incubation in limited glucose media (0.5mM) supplemented with carnitine to promote FAO. During the assay, half the cells are treated with palmitate (C16; a saturated fatty acid) to induce FAO, and half of the C16- and control-treated cells are also treated with etomoxir, an inhibitor of CPT1a, the rate-limiting enzyme of FAO. This allows the contribution of FAO to maintaining the basal metabolic rates of the cells to be assessed. By sequentially injecting the same compounds used in the MST paradigm (oligomycin (0.5µM), FCCP (1µM), rotenone-antimycin A (0.5µM)), we were able to estimate maximal FAO, which provides an estimation of the contribution of FAO to the ability of cells to resolve metabolic stress.

**Figure 5:**
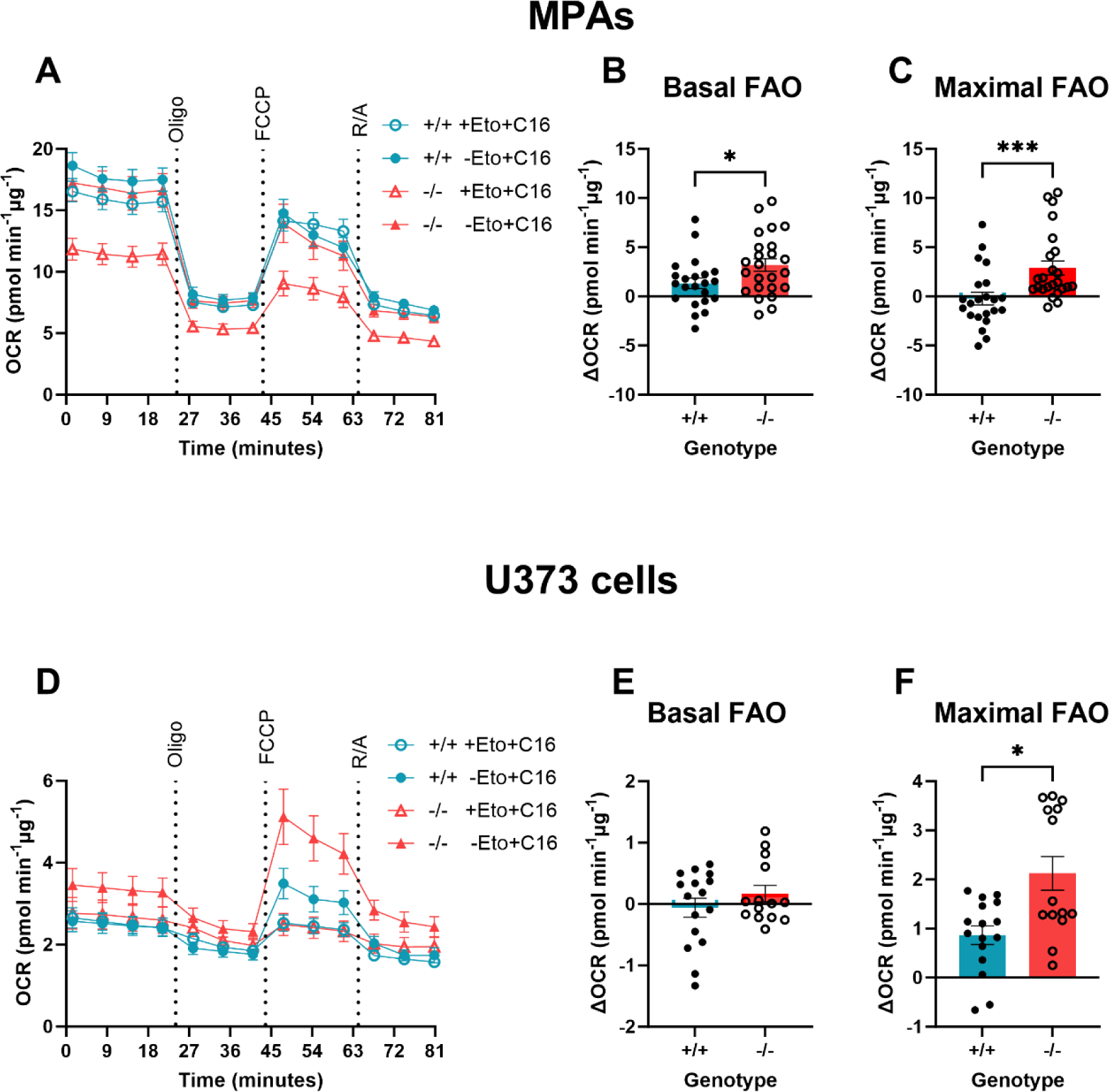
Basal fatty acid oxidation was enhanced in TSPO^-/-^ MPAs. **A:** Oxygen consumption rate (OCR) of TSPO-deficient (TSPO^-/-^) MPAs compared to wildtype (TSPO^+/+^) controls during the fatty acid oxidation stress test. 15 minutes prior to the test, 50% of cells were treated with etomoxir (Eto; 40µM). 50% of cells were treated with palmitate (C16; 200µM) or bovine serum albumin vehicle (BSA; 0.17mM) immediately prior to the test. Oligo = 0.5µM oligomycin, FCCP = 1µM carbonyl cyanide-p-trifluoromethoxyphenylhydrazone, R/A = 0.5µM rotenone/antimycin A. **B:** Basal fatty acid oxidation (FAO) of TSPO^-/-^ MPAs compared to TSPO^+/+^ controls (difference between etomoxir-treated and comparable controls immediately prior to oligo injection). **C**: Maximal FAO of TSPO^-/-^ MPAs compared to TSPO^+/+^ controls (difference between etomoxir-treated and comparable controls after FCCP injection). **D:** Oxygen consumption rate (OCR) of TSPO^-/-^ (KO) U373 cells compared to empty vector (EV) TSPO^+/+^ controls during the fatty acid oxidation stress test. Oligo = 0.5µM oligomycin, FCCP = 1µM carbonyl cyanide-p-trifluoromethoxyphenylhydrazone, R/A = 0.5µM rotenone/antimycin A. **E:** Basal FAO of TSPO^-/-^ U373 cells compared to EV TSPO^+/+^ controls (difference between etomoxir-treated and comparable controls immediately prior to oligo injection). **F**: Maximal FAO of TSPO^-/-^ U373 cells compared to EV TSPO^+/+^ controls (difference between etomoxir-treated and comparable controls after FCCP injection). **A-C**: n=22-24. **D-F:** n-14-16. Data are pooled from across 2 independent plates. Unpaired two-tailed t-test (**B, E**), Mann-Whitney test (**C,F**). *p<0.05, ***p<0.001. Data are expressed as mean ± standard error of the mean.

We found that mean basal FAO was increased by 241.0% in TSPO^-/-^ MPAs (Figure 5B, p=0.031) compared with TSPO^+/+^ MPAs. This suggests that lipids constituted a greater proportion of the substrates used to maintain basal metabolic rates in TSPO^-/-^ MPAs compared with TSPO^+/+^ MPAs. We found that maximal FAO was enhanced in TSPO^-/-^ MPAs (Figure 5C, p=0.0004). In TSPO^-/-^ U373 cells, we did not see any change in basal FAO (Figure 5E, p=0.28), however mean maximal FAO was increased by 245.5% (Figure 5F, p=0.13). Together these data suggest that loss of TSPO increased the contribution of fatty acids as an energy source for MPA metabolism, which may explain how the TSPO^-/-^ MPAs were able to maintain OCR and ECAR in the absence of glucose (Figure 4). The different rates of FAO we observed between U373 astrocytoma cells and MPAs may be due to the relatively increased reliance of MPAs on OCR to meet their bioenergetic requirements (Supplementary Figure 4).

### TSPO formed a complex with CPT1a in U373 cells

Having established that loss of TSPO increases FAO in astrocytes, we began to explore the potential underlying mechanism. Like TSPO, CPT1a resides in the outer mitochondrial membrane^46^. Previous work has established that CPT1a is in a complex with another outer mitochondrial membrane protein, voltage-dependent anion channel (VDAC)^46,47^. VDAC is known to form complexes with TSPO^48^, but the existence of a protein complex containing TSPO and CPT1a has not yet been experimentally confirmed. Our data suggesting that TSPO deficiency increases FAO in astrocytes supports a potential role of TSPO in regulating FAO via a protein complex with CPT1a. To address this, we transiently transfected U373 cells naïve to previous genetic modification with a Myc-DDK tagged TSPO construct. Co-immunoprecipitation was used to identify TSPO-containing complexes via a pull down using anti-Myc bound agarose beads. Using this method, we established that TSPO is part of a protein complex containing both VDAC and CPT1a (Figure 6A, n=4). As this interaction was characterised using transfected cells, to validate this finding we performed a further co-immunoprecipitation using proteins endogenous to U373 cells that were not subject to transfection and determined that this interaction also occurred between endogenous TSPO and CPT1a (Figure 6B,C) (n=3).

**Figure 6:**
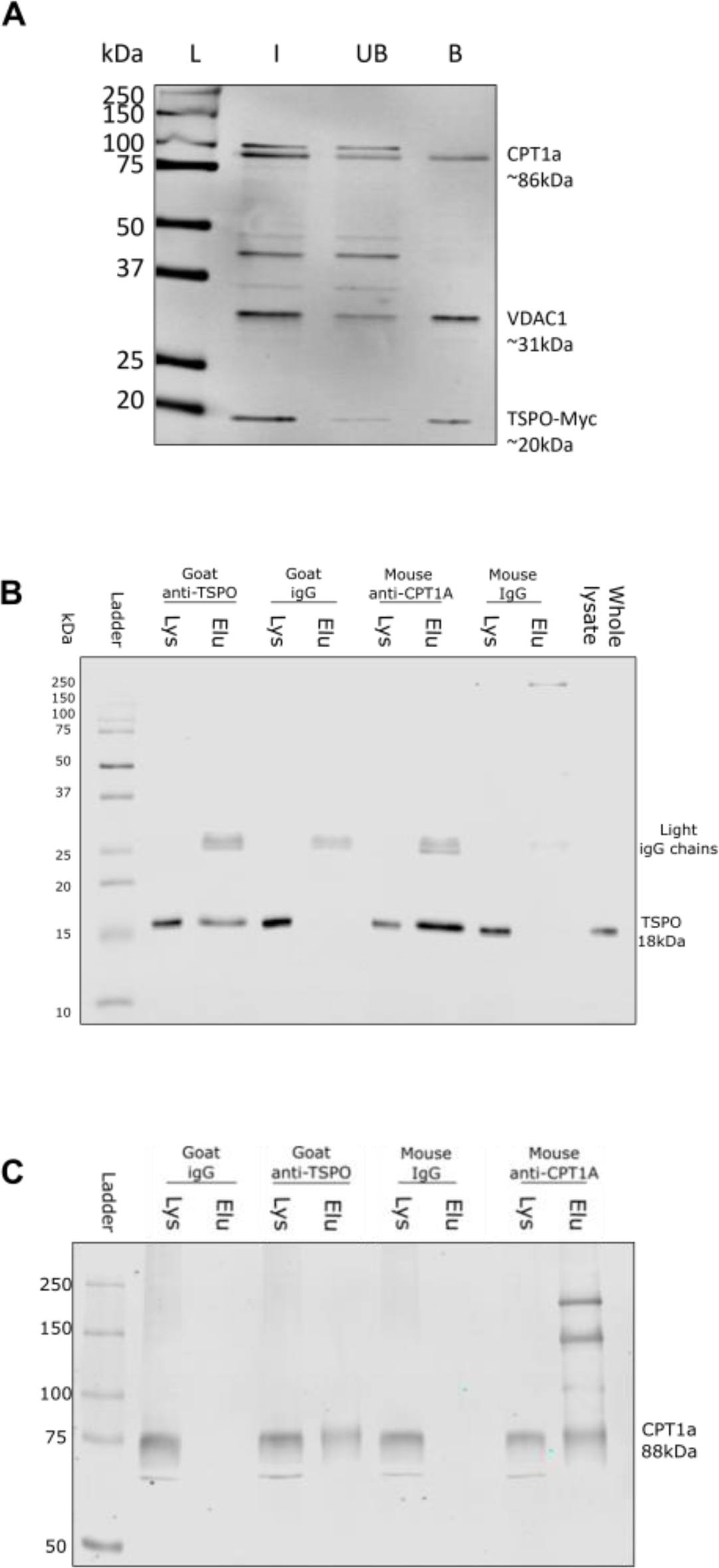
TSPO formed a complex with CPT1a in U373 cells. **A:** Representative immunoblot showing co-immunoprecipitation of a Myc-tagged translocator protein 18kDa (TSPO) construct (TSPO-Myc) with endogenous carnitine palmitoyltransferase 1a (CPT1a) in U373 cells. U373 cells were transiently transfected with TSPO-Myc plasmids (OriGene), and lysate harvested 24h later. Co-immunoprecipitations were performed using Myc-tagged agarose beads (Proteintech). Input (I), unbound fraction (UB) and bound fractions (B) were retained for immunoblotting. Nitrocellulose membranes were blotted for voltage dependent anion channel 1 (VDAC1) (abcam) as a positive control, followed by CPT1a (Proteintech), and Myc (Proteintech) to confirm successful transfection. n=4 separate immunoprecipitations. **B:** Representative immunoblot showing endogenous co-immunoprecipitation of TSPO via CPT1a antibody, with antibody specificity confirmed using species-specific IgG controls. Whole cell lysate was run as a positive control for protein expression, and supernatant (Lys) or eluate (Elu) from the respective antibody (IgG, TSPO, or CPT1a) incubations used to confirm successful co-immunoprecipitation. n=3 separate immunoprecipitations. **C:** Representative immunoblot showing endogenous co-immunoprecipitation of CPT1a via TSPO antibody, with antibody specificity confirmed using species-specific IgG controls. Whole cell lysate was run as a positive control for protein expression, and Lys or Elu from the respective antibody (IgG, TSPO, or CPT1a) incubations used to confirm successful co-immunoprecipitation. n=3 separate immunoprecipitations. L = ladder, kDa = kilodaltons.

### TSPO expression increased in astrocytes after 24h but not 3h LPS stimulation, which temporally corresponded to LPS-induced changes in cellular metabolism

CNS TSPO expression increases during chronic inflammation, such that TSPO is used as a biomarker for monitoring the progression of chronic neuroinflammation^1,7,15,49^. Inflammatory responses incur a metabolic cost, requiring that the energy production of cells increases to maintain a response to inflammatory stimuli^50,51^. We have shown previously that astrocytes undergo a bioenergetic shift to rely more heavily on mitochondrial metabolism after 24h exposure to inflammatory stimuli^28^. Because our data implied that TSPO deficiency alters astrocyte bioenergetics, we hypothesised that altered TSPO expression after 24h LPS stimulation may be an important component of this response. To test this hypothesis, we treated astrocytes isolated from wildtype C57BL/6J neonates with LPS for 3h or 24h, quantified TSPO expression via immunoblotting (Figure 7) and then used a metabolic bioanalyzer to assess the corresponding metabolic changes (Figure 8). The glycolysis inhibitor 2DG was included in Figures 7 and 8 as a positive control for a change in astrocyte metabolic state.

**Figure 7:**
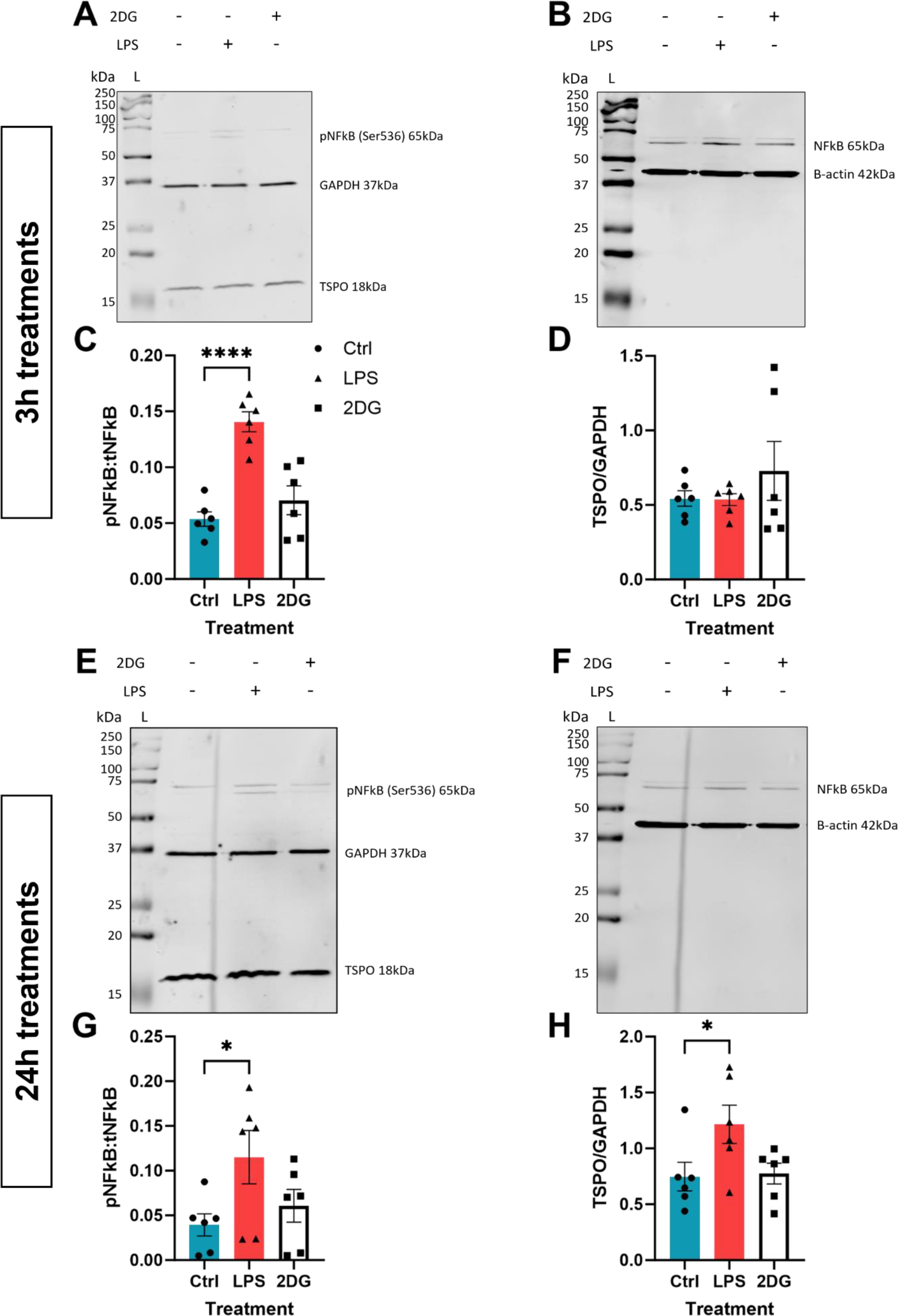
24h but not 3h LPS stimulation increased TSPO expression in C57BL/6J MPAs. **A,B:** Representative immunoblot of C57BL/6J MPA lysate following 3h stimulation with lipopolysaccharide (LPS; 100ng/mL) or 2-deoxyglucose (2DG; 10mM). **A:** Representative immunoblots for phosphorylated nuclear factor kappa B (NFκB) p65 (Ser536) (pNFκB), glyceraldehyde phosphate dehydrogenase (GAPDH), and translocator protein (TSPO). **B:** Representative immunoblots for total NFκB p65 (tNFκB) and β-actin. **C:** Quantification of NFκB activation following 3h LPS or 2DG stimulation expressed as the ratio of pNFκB:NFκB. **D:** Quantification of TSPO expression following 3h LPS or 2DG stimulation. **E,F:** Representative immunoblot of C57BL/6J MPA lysate following 24h stimulation with LPS or 2DG. **E:** Representative immunoblots for pNFκB, GAPDH, and TSPO. **F**: Representative immunoblots for tNFκB and β-actin. **G:** Quantification of NFκB activation following 24h LPS or 2DG stimulation expressed as the ratio of pNFκB:NFκB. **H:** Quantification of TSPO expression following 3h LPS or 2DG stimulation. n=6. One-way ANOVA with Dunnett’s multiple comparisons test (**C,D,E,H**). *p<0.05, ****p<0.0001. Data are expressed as mean ± standard error of the mean.

**Figure 8:**
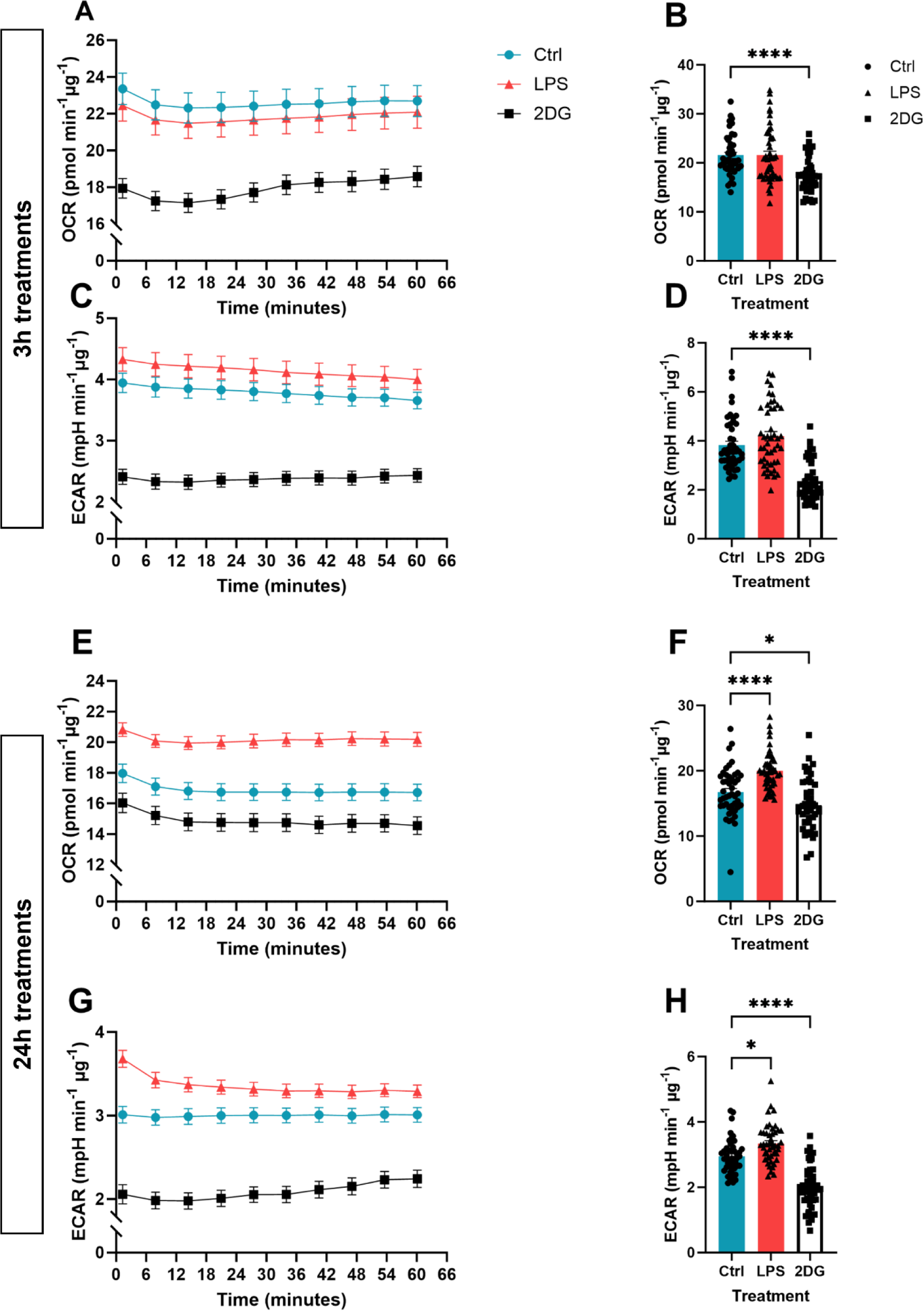
24h LPS stimulation enhanced mitochondrial and non-mitochondrial respiration in C57BL/6J MPAs. **A:** Oxygen consumption rate (OCR) of C57BL/6J MPAs treated with lipopolysaccharide (LPS; 100ng/mL) or 2-deoxyglucose (2DG; 10mM) for 3h. **B:** Quantification of **A**, data are taken from time point 4. **C:** Extracellular acidification rate (ECAR) of C57BL/6J MPAs treated with LPS or 2DG for 3h. **D:** Quantification of **C**, data are taken from time point 4. **E:** OCR of C57BL/6J MPAs treated with LPS or 2DG for 24h. **F:** Quantification of **E**, data are taken from time point 4. **G:** ECAR of C57BL/6J MPAs treated with LPS or 2DG for 24h. **H:** Quantification of **G**, data are taken from time point 4. n=46-48, data are pooled from across 3 independent plates. Ordinary one-way ANOVA with Dunnett’s multiple comparisons test (**B**). Kruskal-Wallis test with Dunn’s multiple comparisons test (**D,F,H**). *p<0.05, ****p<0.0001. Data are expressed as mean ± standard error of the mean.

First, we confirmed that MPAs were undergoing an inflammatory response to LPS stimulation by assessing phosphorylation of nuclear factor kappa-B (p65) which was significantly increased at both timepoints (NFκB; 3h Figure 7A-C; p<0.0001; 24h Figure 7E-G; p=0.044). While 3h LPS stimulation did not significantly increase TSPO expression in MPAs (Figure 7A,D; p=0.99), 24h LPS stimulation induced a significant increase in TSPO expression (Figure 7E,H; p=0.048). Neither 3h nor 24h 2DG stimulation significantly affected NFκB phosphorylation (Figure 7A,D (3h), E,H (24h); p=0.39 (3h), p=0.71 (24h)) or TSPO expression (Figure 7A,D (3h), E,H (24h); p=0.47 (3h), p=0.98 (24h)).

To confirm that the shift in TSPO expression was concomitant with a shift in the metabolic phenotype of MPAs, we next assessed LPS-induced changes in astrocyte metabolism. 3h LPS stimulation did not significantly increase OCR (Figure 8A,B; p=0.99) or ECAR (Figure 8C,D; p=0.59) of MPAs, however, 24h LPS stimulation increased MPA OCR (Figure 8E,F; p<0.0001) and ECAR (Figure 8G,H; p=0.014). 3h 2DG stimulation significantly reduced MPA OCR (Figure 8A,B; p<0.0001) and ECAR (Figure 8C,D; p<0.0001). 24h 2DG stimulation significantly reduced MPA OCR (Figure 8E,F; p=0.048) and ECAR (Figure 8G,H; p<0.0001). While 24h 2DG stimulation significantly reduced cell viability, 24h LPS stimulation did not (Supplementary Figure 5). Though the MPA bioenergetics following 3h LPS stimulation did not match those reported in our previous study^28^, this can be attributed to the current study using different experimental conditions to the former study.

### TSPO deficiency altered the pattern of TNF secretion following LPS stimulation

Having confirmed that the 24h LPS-induced shift in astrocyte metabolic phenotype was concomitant with altered TSPO expression (Figure 7,8), we postulated that TSPO-deficient astrocytes may show altered immunometabolic responses to LPS. We began by characterising secretion of TNF from TSPO^-/-^ MPAs following LPS stimulation (Figure 9). Following 3h LPS stimulation, we observed a statistically significant effect of genotype (Figure 9A; p_genotype_<0.0001, F_(1,16)_ = 141.2), LPS stimulation (p_LPS_=0.0007, F_(1,16)_ = 17.41) and an interaction (p_interaction_=0.0007, F_(1,16)_ = 17.41) of these variables on TNF secretion. Likewise, following 24h LPS stimulation, we observed a statistically significant effect of genotype (Figure 9B; p_genotype_=0.01, F_(1,20)_ = 8.038), LPS stimulation (p_LPS_<0.0001, F_(1,20)_ = 91.97) and an interaction (p_interaction_=0.01, F_(1,20)_ 8.038) of these variables on TNF secretion. Post-hoc analyses confirmed that in response to 3h LPS stimulation, TNF secretion from TSPO^-/-^ MPAs was reduced by 52.0% compared to TSPO^+/+^ controls (Figure 9A; p=0.0001). In contrast, after 24h LPS stimulation, we found that TSPO^-/-^ MPAs released 183.9% more TNF than TSPO^+/+^ controls (Figure 9B, p=0.0035).

**Figure 9:**
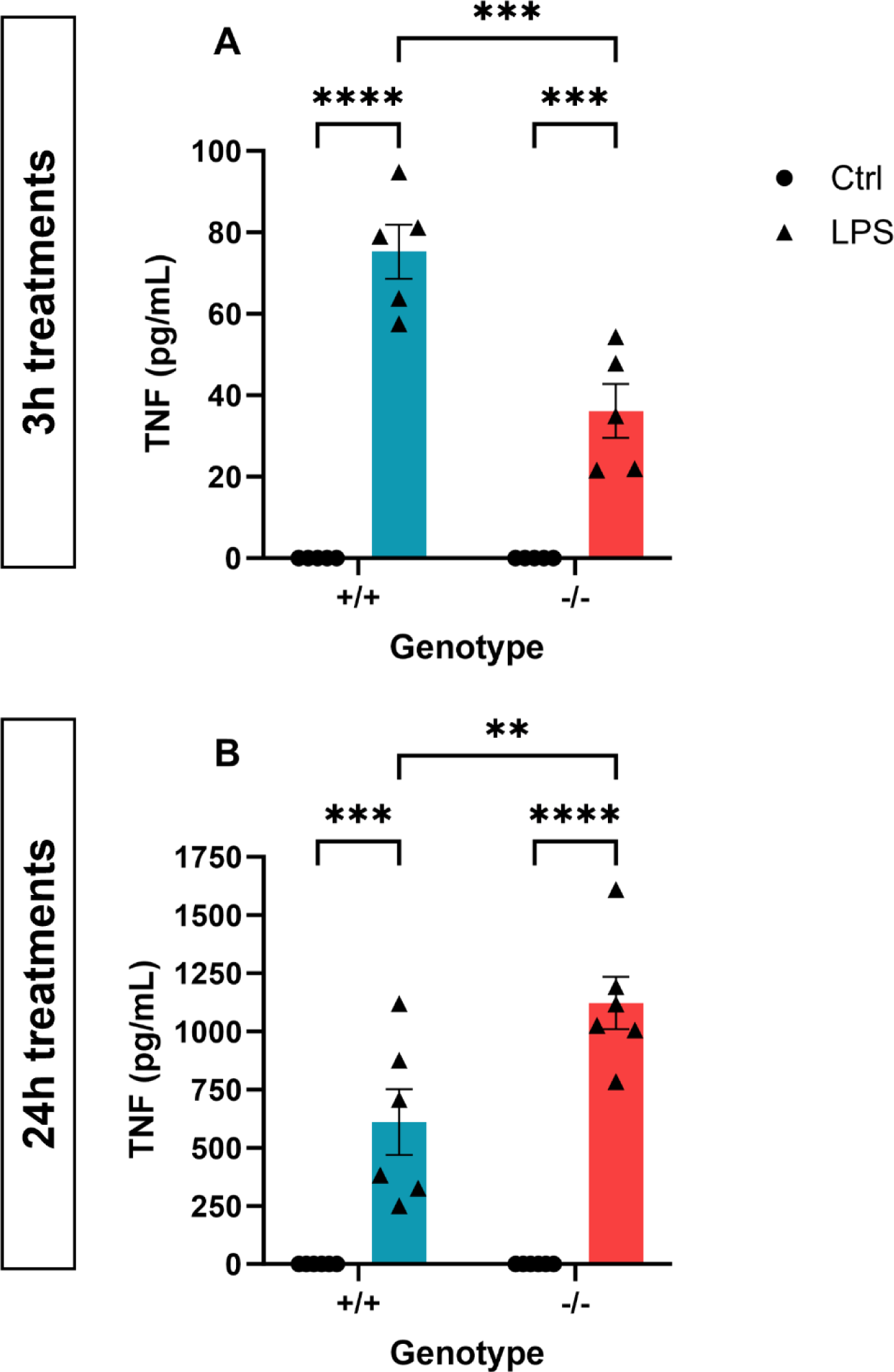
TSPO deficiency modulated cytokine release from MPAs. **A,B:** Tumour necrosis factor (TNF) secretion from TSPO-deficient (TSPO^-/-^) MPAs and wildtype (TSPO^+/+^) controls following 3h (**A**) or 24h (**B**) lipopolysaccharide (LPS; 100ng/mL) stimulation. 2-way ANOVA with Šídák’s multiple comparisons test. **A:** p_genotype_<0.0001, F_(1,16)_ = 141.2. p_LPS_=0.0007, F_(1,16)_ = 17.41. p_interaction_=0.0007, F_(1,16)_ = 17.41. **B:** p_genotype_=0.0102, F_(1,20)_ = 8.038. p_LPS_<0.0001, F_(1,20)_ = 91.97. p_interaction_=0.0102, F_(1,20)_ 8.038. n=6. **p<0.01, ***p<0.001, ****p<0.0001. Data are expressed as mean ± standard error of the mean.

As a key regulator of the inflammatory response, altered NFκB phosphorylation in TSPO^-/-^ MPAs may have contributed to this. However, we observed no statistically significant effect of genotype (Figure 10A-C; p_genotype_=0.3408, F_(1,19)_ = 0.95) on NFκB phosphorylation following 3h LPS stimulation, but observed a significant effect of LPS stimulation on NFκB phosphorylation following 3h stimulation (p_LPS_<0.0001, F_(1,19)_ = 41.95). We observed no statistically significant interaction between TSPO genotype and 3h LPS stimulation (p_interaction_=0.18, F_(1,19)_ = 1.911). Likewise, following 24h LPS stimulation, we observed no statistically significant effect of genotype (Figure 10D-F; p_genotype_=0.97, F_(1,19)_ = 0.001168). However, we observed a statistically significant effect of 24h LPS stimulation on NFκB phosphorylation (p_LPS_<0.0001, F_(1,19)_ = 25.52). There was no statistically significant interaction between these variables (p_interaction_=0.57, F_(1,19)_ = 0.3276). Post-hoc assessment confirmed no significant difference in NFκB phosphorylation in TSPO^-/-^ MPAs compared to TSPO^+/+^ controls following either 3h (Figure 10A-C; p=0.48) or 24h LPS stimulation (Figure 10D-F; p=0.99).

**Figure 10:**
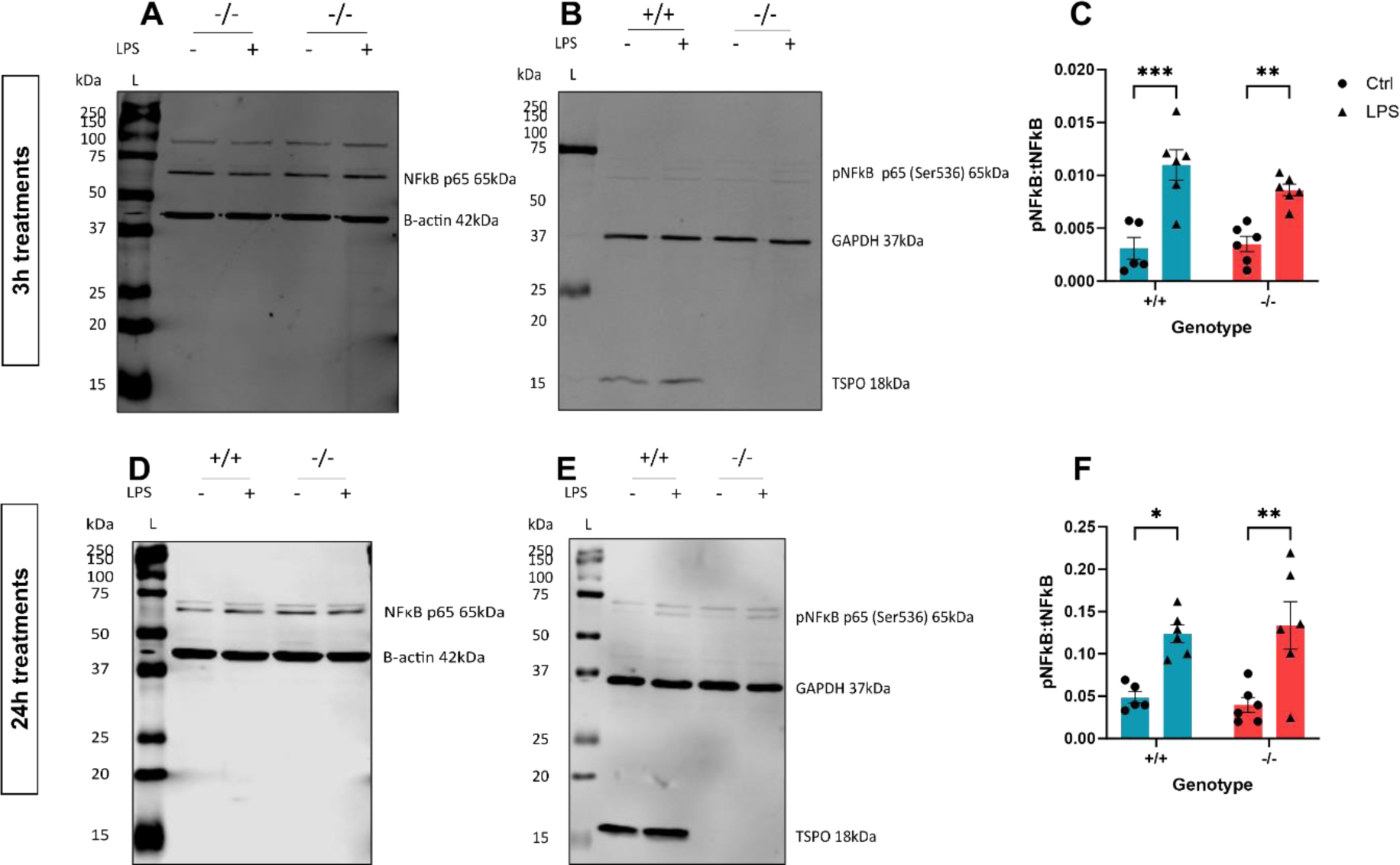
TSPO deficiency did not modulate NFκB activation following LPS stimulation. **A,B**: Representative immunoblot of cell lysate from Figure 9A. **A:** Representative immunoblot for total nuclear factor kappa B p65 (NFκB, tNFκB) and β-actin (B-actin) 3h ± lipopolysaccharide (LPS; 100ng/mL) stimulation. **B:** Representative immunoblot for phosphorylated NFκB p65 (Ser536; pNFκB), glyceraldehyde phosphate dehydrogenase (GAPDH), translocator protein 18kDa (TSPO) 3h ± LPS stimulation. **C:** Quantification of NFκB activation following 3h ± LPS stimulation expressed as the ratio of pNFκB:NFκB. **D,E**: Representative immunoblot of cell lysate from Figure 9B. **D:** Representative immunoblot for tNFκB and β-actin 24h ± LPS stimulation. **E:** Representative immunoblot for pNFκB (Ser536), GAPDH, and TSPO 24h ± LPS stimulation. **F:** Quantification of NFκB activation following 24h ± LPS stimulation expressed as the ratio of pNFκB:NFκB. **C,F:** 2-way ANOVA with Šídák’s multiple comparisons test. **C:** p_genotype_=0.3408, F_(1,19)_ = 0.9547. p_LPS_<0.0001, F_(1,19)_ = 41.95. p_interaction_=0.1829, F_(1,19)_ = 1.911. **F:** p_genotype_=0.9731, F_(1,19)_ = 0.001168. p_LPS_<0.0001, F_(1,19)_ = 25.52. p_interaction_=0.5738, F_(1,19)_ = 0.3276. n=6. *p<0.05, **p<0.01, ***p<0.001. Data are expressed as mean ± standard error of the mean.

### Astrocyte TSPO deficiency did not impact the metabolic response following LPS stimulation

We hypothesised that the altered cytokine secretion profile observed in TSPO^-/-^ MPAs might reflect a different metabolic response to LPS in these cells, related to their altered metabolic profile (Figures 1-5).

Following 3h LPS stimulation, the statistically significant impact of genotype on astrocyte metabolic parameters seen in earlier experiments (Figure 1) was maintained (Figure 11A-D; OCR: p_genotype_<0.0001, F_(1,66)_ = 18.45; ECAR: p_genotype_=0.0019, F_(1,64)_ = 10.51). However, using a two-way ANOVA we observed no statistically significant effect of 3h LPS stimulation on MPA bioenergetics (OCR p_LPS_=0.54, F_(1,66)_ = 0.3802; ECAR p_LPS_=0.72, F_(1,20)_ = 0.1259), and observed no statistically significant interaction between genotype and 3h LPS stimulation (Figure 11A-D; OCR p_interaction_=0.26, F_(1,66)_ = 1.3002, ECAR p_interaction_=0.64, F_(1,64)_ = 0.2173). Post-hoc analysis revealed that, in line with our previous data from C57BL/6J MPAs (Figure 7), neither the OCR nor ECAR of TSPO^+/+^ MPAs significantly changed in response to 3h LPS stimulation (Figure 11A-D; p=0.66 [OCR], p=0.99 [ECAR]).

**Figure 11:**
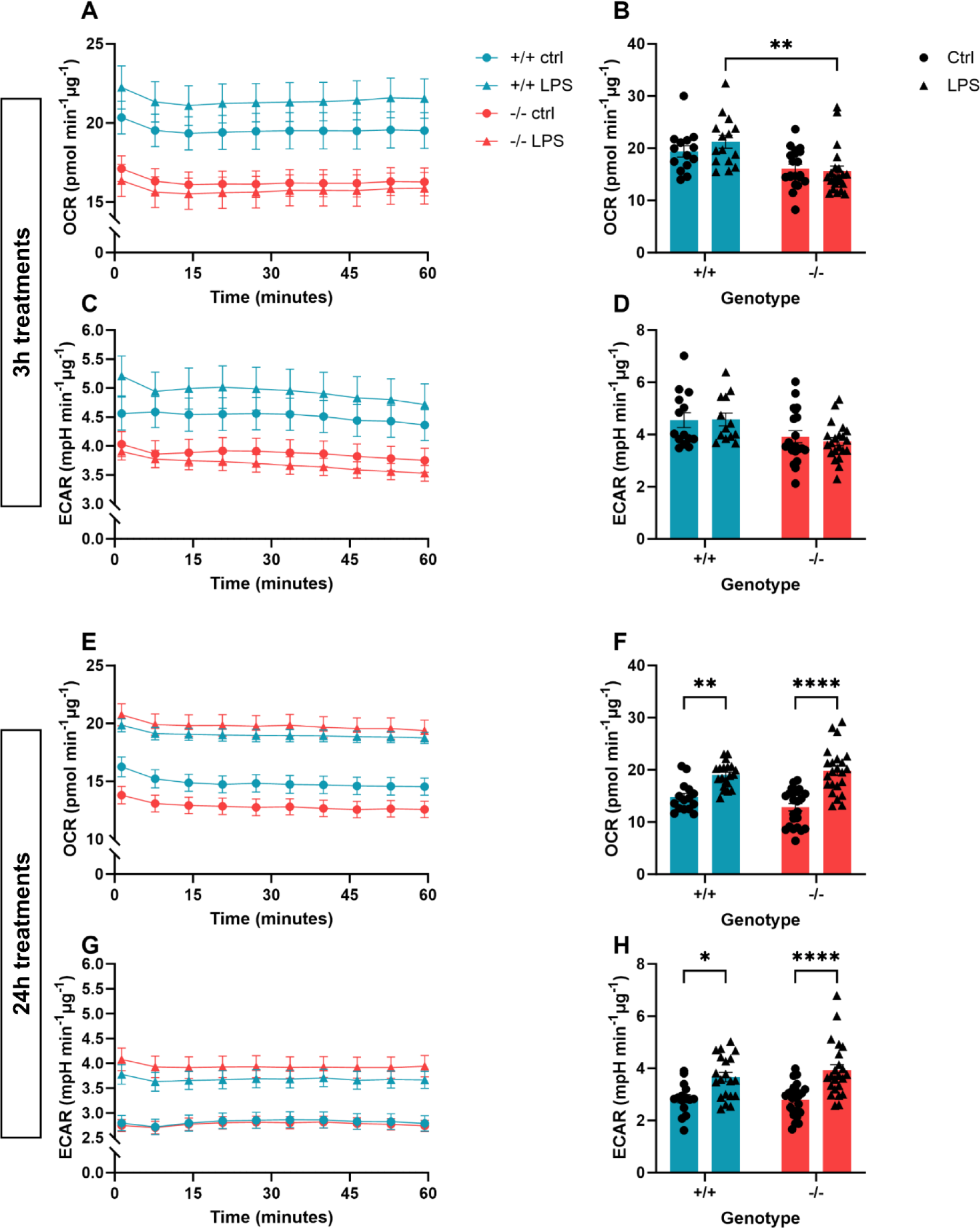
TSPO deficiency did not affect the metabolic response of MPAs to inflammatory stimulation. **A:** Oxygen consumption rate (OCR) of TSPO-deficient (TSPO^-/-^) MPAs and wildtype (TSPO^+/+^) controls 3h ± lipopolysaccharide (LPS; 100ng/mL) stimulation. **B:** Quantification of **A**, data were taken from time point 4. **C:** Extracellular acidification rate (ECAR) of TSPO^-/-^ MPAs and TSPO^+/+^ controls 3h ± LPS stimulation. **D:** Quantification of **C**, data were taken from timepoint 4. **E:** OCR of TSPO^-/-^ MPAs and TSPO^+/+^ controls 24h ± LPS stimulation. **F:** Quantification of **E**, data were taken from timepoint 4. **G**: ECAR of TSPO^-/-^ MPAs and TSPO^+/+^ controls 24h ± LPS stimulation. **H:** Quantification of **G**, data were taken from timepoint 4. **B,D,F,H:** 2-way ANOVA with Tukey’s multiple comparisons test. **B:** p_genotype_<0.0001, F_(1,66)_ = 18.45. p_LPS_=0.5396, F_(1,66)_ = 0.3802. p_interaction_=0.2579, F_(1,66)_ = 1.3002. **D**: p_genotype_=0.0019, F_(1,64)_ = 10.51. p_LPS_=0.7230, F_(1,20)_ = 0.1259. p_interaction_=0.6427, F_(1,64)_ = 0.2173. **F:** p_genotype_=0.4897, F_(1,78)_ = 0.4818. p_LPS_<0.0001, F_(1,78)_ = 52.35. p_interaction_=0.0844, F_(1,78)_ = 3.055. **H**: p_genotype_=0.5445, F_(1,78)_ = 0.3704. p_LPS_<0.0001, F_(1,78)_ = 28.89. p_interaction_=0.4141, F_(1,78)_ = 0.6742. **B,D:** n=15-21 per group, data are pooled from across 2 independent plates. **F,H:** n=15-24 per group, data are pooled from across 2 independent plates. *p<0.05, **p<0.01, ****p<0.0001.

Following our 24h LPS stimulation paradigm overall there was no statistically significant impact of TSPO genotype on MPA OCR (Figure 11E-H; p_genotype_=0.49, F_(1,78)_ = 0.4818) or ECAR (p_genotype_=0.54, F_(1,78)_ = 0.3704). LPS stimulation significantly impacted both OCR (p_LPS_<0.0001, F_(1,78)_ = 52.35) and ECAR (p_LPS_<0.0001, F_(1,78)_ = 28.89), but there was not a statistically significant interaction between genotype and 24h LPS stimulation (OCR: p_interaction_=0.084, F_(1,78)_ = 3.055; ECAR: p_interaction_=0.41, F_(1,78)_ = 0.6742). However, post-hoc analyses revealed that, in response to 24h LPS stimulation both OCR and ECAR increased in TSPO^+/+^ (Figure 11F, 29.0% [OCR], p=0.003; Figure 11H, 29.1% [ECAR], p=0.02) and TSPO^-/-^ MPAs (Figure 11F, 54.4% [OCR], p<0.0001; Figure 11H 40.1% [ECAR], p<0.0001). Despite this, there was no statistically significant difference in the metabolic responses of TSPO^-/-^ MPAs to 24h LPS stimulation compared to TSPO^+/+^ controls (Figure 11F,H; p=0.87 [OCR], p=0.74 [ECAR]).

We also assessed changes in expression of glucose transporter 1 (GLUT1), which we have previously shown to be modulated by 24h but not 3h LPS stimulation in MPAs^28^. We observed neither a statistically significant impact of TSPO genotype (Figure 12A,B p=_genotype_0.87, F_(1,19)_ = 0.027) nor of 3h LPS stimulation on GLUT1 expression (p_LPS_=0.63, F_(1,19)_ = 0.24). We did not observe a statistically significant interaction between 3h LPS stimulation and TSPO genotype on GLUT1 expression (p_interaction_=0.17, F_(1,19)_ = 2.0). Post-hoc analyses found no significant effect of genotype on GLUT1 expression at baseline or following 3h LPS stimulation in these cells, which showed the same response as TSPO^+/+^ controls (Figure 12A,B; p=0.96).

**Figure 12:**
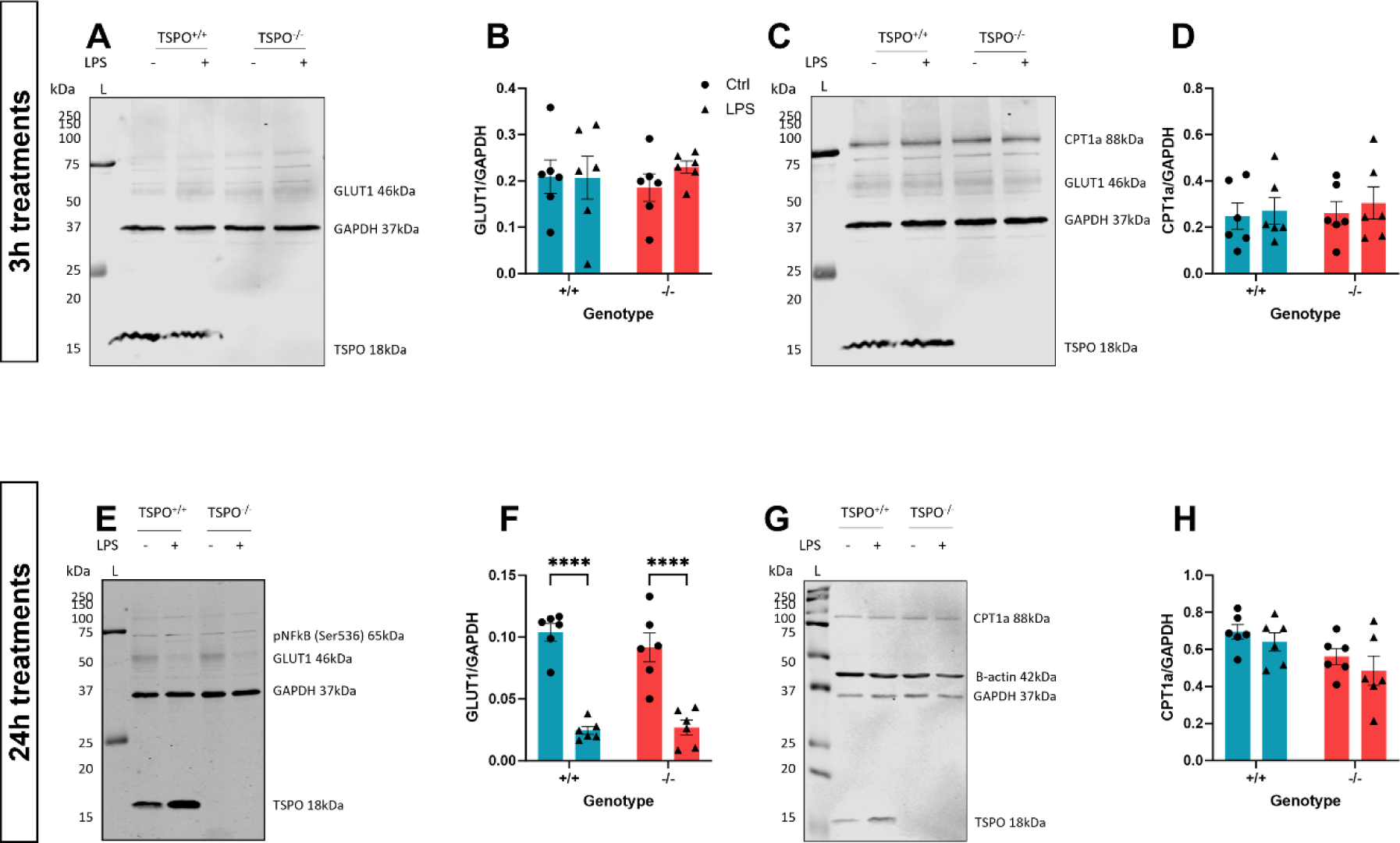
Inflammation-induced changes to GLUT1 and CPT1a expression were not modulated by TSPO deficiency. **A:** Representative immunoblot of cell lysate from Figure 9A for GLUT1 following 3h LPS stimulation. **B:** Quantification of **A**. **C:** Representative immunoblot of cell lysate from Figure 9A for CPT1a following 3h LPS stimulation. **D:** Quantification of **C**. **E:** Representative immunoblot of cell lysate from Figure 9B for GLUT1 following 24h LPS stimulation. **F:** Quantification of **E**. **G:** Representative immunoblot of cell lysate from Figure 9B for CPT1a following 24h LPS stimulation. **H:** Quantification of **G**. **B,D,F,H:** 2-way ANOVA with Šídák’s multiple comparisons test. **B:** p_genotype_=0.9904, F_(1,20)_ = 0.0001496. p_LPS_=0.5423, F_(1,20)_ = 0.3843. p_interaction_=0.4914, F_(1,20)_ = 0.4914. **D:** p_genotype_=0.6950, F_(1,20)_ = 0.1582. p_LPS_=0.5721, F_(1,20)_ = 0.3300. p_interaction_=0.8628, F_(1,20)_ = 0.03066. **F:** p_genotype_=0.5249, F_(1,20)_ = 0.4188. p_LPS_<0.0001, F_(1,20)_ = 89.35. p_interaction_=0.3435, F_(1,20)_ = 0.9413. **H:** p_genotype_=0.0165, F_(1,20)_ = 6.848. p_LPS_=0.2485, F_(1,20)_ = 1.413. p_interaction_=0.8470, F_(1,20)_ = 0.03820. n=6. ****p<0.0001. Data are expressed as mean ± standard error of the mean.

In contrast to our expectations, following 24h LPS stimulation, TSPO genotype had no statistically significant impact on GLUT1 expression (Figure 12E,F; p_genotype_=0.52, F_(1,20)_ = 0.42). In line with our previous study^28^, 24h LPS stimulation had a statistically significant impact on GLUT1 expression (Figure 12E,F; p_LPS_<0.0001, F_(1,20)_ = 89.35), but we observed no statistically significant interaction between TSPO genotype and LPS stimulation on GLUT1 expression (p_interaction_=0.34, F_(1,20)_ = 0.91). Using post-hoc analyses, we found that in line with wildtype controls, GLUT1 expression was similarly reduced by 24h LPS stimulation in MPAs (Figure 12E,F; p<0.0001) regardless of TSPO-genotype (Figure 12E,F; p>0.99).

Previous work examining the role of TSPO in cellular metabolism has linked TSPO deficiency to increased expression of the *Cpt1a* gene in MA-10 Leydig cells^31^, however, to the best of our knowledge, complementary changes to the expression of the CPT1a protein in astrocytes following genetic ablation of TSPO have not been evaluated. We observed no statistically significant effect of TSPO genotype (Figure 12C,D, p_genotype_=0.70, F_(1,20)_ = 0.16) or 3h LPS stimulation (p_LPS_=0.57, F_(1,20)_ = 0.33) on CPT1a expression, and observed no statistically significant interaction between these two variables (p_interaction_=0.86, F_(1,20)_ = 0.0031). In contrast, using our 24h LPS stimulation model, we observed a statistically significant effect of genotype on CPT1a expression (Figure 12 G,H; p_genotype_=0.017, F_(1,20)_ = 6.8). However, we observed no statistically significant effect of 24h LPS stimulation on CPT1a expression (p_LPS_=0.25, F_(1,20)_ = 1.4), nor did we observe an interaction between TSPO genotype and 24h LPS stimulation (p_interaction_=0.85, F_(1,20)_ = 0.038). Post-hoc analysis confirmed that CPT1a expression was unchanged by TSPO genotype at baseline or following 3h (Figure 12E,F; p=0.99 (TSPO^+/+^), p=0.95 (TSPO^-/-^)) or 24h LPS stimulation (Figure 12G,H; p=0.89 (TSPO^+/+^), p=0.76 (TSPO^-/-^)).

## Discussion

TSPO, a protein linked to a variety of cellular processes including regulation of cellular metabolism and inflammatory responses, has previously been demonstrated to play a role in fatty acid metabolism in Leydig cells^31^, adrenal glands^52^, and hepatocytes^53^. However, it is unclear whether this happens in all cell types and by what underlying mechanisms TSPO may be involved with FAO. In this study, we provide data supporting the hypothesis that TSPO regulates cellular bioenergetics in astrocytes. We showed that TSPO^-/-^ astrocytes secrete less lactate than wildtype controls and rely more on FAO to meet their bioenergetic requirements. For the first time, we demonstrated using co-immunoprecipitation an interaction between TSPO and the rate-limiting enzyme in FAO, CPT1a, providing a possible mechanism through which TSPO may regulate FAO. We showed here that loss of TSPO expression does not restrict the metabolic response of astrocytes to an inflammatory stimulus, nor alter the expression of key metabolic proteins in response to inflammation, but may alter the temporal profile of the astrocyte inflammatory response.

### TSPO as a regulator of astrocyte metabolism

In other glial and non-glial cells, TSPO deficiency has been linked to reduced oxidative phosphorylation and glycolysis^9,30,31,54,55^. In line with these studies, we found that TSPO deficiency in astrocytes reduced basal mitochondrial metabolism and non-mitochondrial respiration. Furthermore, in agreement with a recent study in microglia^9^, we found that TSPO deficiency enhances the metabolic response in astrocytes following the injection of a supraphysiological glucose concentration during the glycolysis stress test. This may be interpreted as increased glycolytic rate in TSPO^-/-^ MPAs. During a nutrient deficit simulated by exposing the cells to a glucose-free condition for 1h, we found that TSPO-deficient astrocytes lacked the metabolic adaptations observed in TSPO-expressing controls. Moreover, we found that the bioenergetic rates of TSPO^-/-^ astrocytes did not change significantly between glucose and glucose-free conditions, and TSPO^-/-^ astrocytes secreted significantly less L-lactate in the presence of glucose. This contrasts with a recent publication demonstrating that TSPO deficiency in microglia increases lactate production and glycolysis^55^. The difference between these findings may be due to fundamental differences in the metabolic machinery of astrocytes (which express relatively high amounts of CPT1a^35^) and microglia (which express less CPT1a than astrocytes^34,35^) underlying the resulting metabolic profile. Regardless, our data show that the basal bioenergetic rates of TSPO^-/-^ astrocytes are reduced relative to TSPO^+/+^ controls. This may be because TSPO^-/-^ astrocytes metabolise less glucose to maintain their bioenergetic profiles, or that in these cells more products of glycolysis undergo oxidative phosphorylation.

As we did not directly quantify pyruvate production in these cells our L-lactate data alone cannot inform us whether TSPO^-/-^ MPAs oxidise a greater amount of lactate than wildtype controls. Because mitochondrial pyruvate metabolism consumes oxygen, one may therefore anticipate that enhanced mitochondrial pyruvate metabolism would elicit a concomitant increase in OCR. However, when exogenous glucose was unavailable as a substrate, we saw no significant increase in the OCR of TSPO^-/-^ astrocytes (Figure 4). Thus, we are confident that the decrease in L-lactate secreted by TSPO^-/-^ astrocytes relative to TSPO^+/+^ controls indicates that TSPO^-/-^ astrocytes are less dependent on glucose to meet their bioenergetic requirements. Furthermore, removal of glucose from the media resulted in no change in lactate secretion, OCR or ECAR in TSPO^-/-^ astrocytes. This suggests that loss of TSPO may enhance the metabolic flexibility of astrocytes, potentially facilitating metabolism of other fuel sources even when glucose is available as a metabolic substrate. Therefore, we speculated that TSPO^-/-^ astrocytes may have been better able to utilise alternative metabolic substrates.

### Regulation of FAO in astrocytes by TSPO

TSPO has previously been linked to the regulation of fatty acid oxidation (FAO)^31,52,53^. We postulated that FAO made a larger contribution to maintaining basal metabolic rates in TSPO^-/-^ astrocytes, as has been reported in Leydig cells^31^ and hepatocytes^53^. Using the FAO stress test paradigm, we found that TSPO^-/-^ astrocytes indeed utilised more fatty acids to maintain their basal bioenergetic needs: OCR was increased in TSPO^-/-^ MPAs supplemented with palmitate compared to those supplemented with palmitate and treated with etomoxir, a potent inhibitor of CPT1a, implying that FAO formed a larger part of the metabolic base of TSPO^-/-^ MPAs compared to wildtype MPAs (Figure 5B).

While the role of TSPO in regulating FAO has been reported before^31,52,53^, a mechanism through which it is driven has only been speculated upon with no direct experimental confirmation to our knowledge. TSPO is reported to have multiple interacting partners, including VDAC^2,56^, another constituent of the outer mitochondrial membrane. By transiently inducing overexpression of a tagged TSPO construct in our astrocytoma lines, we were able to demonstrate that the tagged variant not only interacts with VDAC (Figure 6A) – as predicted – but also with CPT1a. We were then able to recapitulate this finding in U373 cells using endogenous TSPO (Figure 6B,C), demonstrating that this interaction was not a result of genetically manipulating these cells. Unfortunately, we were unable to successfully repeat this using MPAs due to technical issues related to the cellular biomass required. Regardless, given that loss of TSPO has previously been linked to increased FAO, and overexpression of TSPO in similar cell lines has reduced expression of *Cpt1a* genes^31^, this may begin to shed light on the underlying role of the TSPO-VDAC-CPT1a complex in astrocytes *in vitro* and *in vivo*. We propose that this interaction is a mechanism by which TSPO regulates FAO in astrocytes. Overall, our data interrogating the role of TSPO in regulating astrocyte metabolism suggests that TSPO deficiency promotes FAO. This implies that, when present, TSPO may act to suppress FAO in these cells, potentially via its interaction with CPT1a.

Fairley *et al*.^55^ recently observed that TSPO also forms a complex with HK2, the rate-limiting enzyme of glycolysis. Because microglia express low or no levels of CPT1a^34,35^, a TSPO-CPT1a interaction is less likely to be detected/present in these cells. However, both microglia^55^ and astrocytes express HK2^57^, therefore it is likely that the TSPO-HK2 interaction may also be conserved in astrocytes. While we did not explore a potential interaction of TSPO with HK2 in astrocytes, this remains an intriguing avenue for future study. TSPO and HK2 have both been reported to interact with VDAC^2,56,58^, thus existence of a dynamic TSPO-HK2-CPT1a-VDAC multimer in astrocytes remains plausible. Because metabolic pathways do not exist in isolation, and multiple pathways are used simultaneously to meet the cellular bioenergetic requirements it is plausible that TSPO acts at the interface of CPT1a and HK2 to regulate the balance between glycolysis and FAO.

### Loss of TSPO and the astrocyte metabolic response to inflammation

In a pathophysiological context, increased use of FAO has been linked to an anti-inflammatory (‘A2’)-like phenotype in astrocytes^51^. In our data, when mitochondrial stress responses were assessed during the FAO paradigm using the uncoupling agent FCCP – to mimic the oxidative stress that might be seen in disease states – we found that TSPO^-/-^ astrocytes used more FAO to fuel their response (Figure 5C,F). This suggests that the bioenergetic response of TSPO^-/-^ MPAs to inflammatory stimulation may be modulated by enhanced FAO. However, we found that the bioenergetic phenotype of LPS stimulated TSPO^-/-^ MPAs was not significantly different from TSPO^+/+^ controls (Figure 11), suggesting that the basal metabolic response of these cells to inflammatory stimuli is not modulated by the absence of TSPO, at least using LPS as an inflammatory stimulus. We did not measure FAO following LPS stimulation, so we cannot provide further comment on whether enhanced FAO in TSPO^-/-^ MPAs modulates the bioenergetic response. Despite this, we have presented data suggesting an altered temporal release profile of TNF from TSPO^-/-^ MPAs (Figure 9), though we only examined one concentration of one inflammatory stimulus at two timepoints. A cytokine array panel may provide valuable insight into potential modulatory effects of TSPO expression on the release of a wider range of cytokines. This could be paired with a greater variety of inflammatory stimuli, concentrations and timepoints to provide a more holistic insight. The underlying mechanism may be addressed by interrogating Ca^2+^ dynamics: TSPO has been linked to calcium signalling in mouse embryonic fibroblasts and neurons^59^ *in vitro*, and tanycytes *in vivo*^8^, which may hint at a potential underlying mechanism distinct from a shift in the metabolic phenotype in astrocytes. Future studies could further interrogate the mechanism by which TSPO regulates the reactive phenotype of astrocytes, perhaps via transcriptomic arrays, which we did not examine in this study so cannot provide comment on. This could be paired with a metabolomics array to better elucidate any differences in substrate utilisation following ‘acute’ and ‘chronic’ proinflammatory stimulation of TSPO^-/-^ astrocytes.

Although it is appreciated that pharmacological inhibition of a protein may have different impacts than genetic deletion, our data demonstrating a role for TSPO in regulating astrocyte metabolism and potentially impacting the release of TNF from astrocytes may have implications when considering glial TSPO as a therapeutic target. Firstly, astrocytes are commonly thought to ‘feed’ neurons lactate for downstream metabolism as part of the astrocyte-neuron lactate shuttle^60^, and we have reported that TSPO^-/-^ astrocytes produce less lactate (Figure 4), implying that these astrocytes may be less able to provide trophic support to neurons as has been speculated elsewhere^61^. Thus, we can speculate that prolonged pharmacological inhibition of astrocyte TSPO may result in an impaired ability of astrocytes to sufficiently support neuronal metabolism for prolonged periods. Given recent evidence for the role of astrocytes in regulating memory formation^62^ and other behaviours^63–66^, and the crucial role of astrocyte CPT1a in cognition^67^, prolonged inhibition of astrocyte TSPO may yield detrimental effects on cognitive and other neural functions which may need to be further investigated. Secondly, astrocytes are highly plastic cells intrinsically involved with the maintenance of the brain microenvironment^19,68^. Perturbations to the ability of astrocytes to respond to noxious stimuli, including by modulating their metabolism – such as chronic neuroinflammation^19,22^ – may in turn impair the ability of these cells to resolve inflammation and restore homeostasis. While this impaired metabolic response to LPS-induced inflammation was not observed in our studies, we only looked at one inflammatory stimulus at one concentration, so additional comprehensive studies are required. TSPO may serve as an attractive therapeutic target due to the proposed anti-inflammatory effects of its ligands^36,69–73^. If TSPO indeed functions as a regulator of metabolic flexibility in astrocytes, continuous pharmacological inhibition of TSPO function in these cells may blunt their ability to respond to inflammatory and other stimuli when chronically administered. However, a better understanding of the physiology and pharmacology of this protein is required.

### Conclusion

We have shown that loss of TSPO reduced the basal metabolic rates of astrocytes, and enabled astrocytes to maintain their metabolic rates in the absence of glucose while reducing L-lactate secretion. Together these data suggest that TSPO deficient astrocytes are less reliant on glucose to meet their energetic needs. Crucially, we have shown that the rate of FAO is increased in TSPO^-/-^ astrocytes and that, when present, TSPO forms a complex with CPT1a, reinforcing this concept. This suggests that TSPO may ordinarily inhibit CPT1a function to conserve intracellular fat stores. Taken together, these data show that TSPO may act as a regulator of metabolic flexibility in astrocytes. Finally, we have shown that despite increases in basal FAO, the metabolic response to LPS stimulation is not impeded in TSPO^-/-^ astrocytes, though the release profile of the pro-inflammatory cytokine TNF is altered. Recapitulation of our results via pharmacological inhibition of TSPO may be important to decipher the therapeutic potential of this protein.

## Acknowledgements

This work was supported by grants from the British Society for Neuroendocrinology (WF) and Research England – Expanding Excellence in Diabetes award, and internal funding from the University of Exeter Medical School (which supported JLR). The TSPO-deficient mice were generated from material provided by the European Mouse Mutant Cell Repository through funding from the Medical Research Council, Grant/Award Number: MR/R014345/1 (to KLJE and CB).

The authors would like to extend their thanks to the Exeter BRF for facilitating and providing assistance with our animal work, and to Dr Ana Miguel Cruz, Dr Katie Partridge, Dr Paul Weightman Potter, and Dr Jiping Zhang for their thoughts and insights.

