## supplementary for "Regulation of astrocyte metabolism by mitochondrial translocator protein 18kDa"

### Supplementary Materials

#### Supplementary Table 1: CRISPR-Cas9 gRNA sequences

| gRNA name: | Target exon: | Sequence (5'-3'): |
|---|---|---|
| Ex2a | 2 | GAGGGTCTCCGCTGGTACGC |
| Ex2b | 2 | TCTGCAGGCCGGCGTACCAG |
| Ex3a | 3 | CGGGCACCAAAGAAGATGGG |
| Ex3b | 3 | TCGGGCACCAAAGAAGATGG |

#### Supplementary Table 2: Primers used to genotype CRISPR-Cas9-generated TSPO-KO and EV U373 cells

| Product: | Forward primer (5'-3'): | Reverse primer (5'-3'): | Product size (bp): |
|---|---|---|---|
| *Tspo* exon 2 | GGAAGAGAGGAGGGGTCAGT | CAGGCGTTCAACAAAGGAGC | 821 |
| *Tspo* exon 3 | CAGGCCAAATGTCCCATCCT | ACAGTACCAGGCTTCGTGTG | 751 |
| *Tspo* exons 2 and 3 | CTGGGACGAAAGTGCTCAGT | ACACAGTACCAGGCTTCGTG | 3253 |

#### Supplementary Table 3: Modified RIPA buffer

| Compound: | Concentration (mM): | Catalogue number: | Supplier: |
|---|---|---|---|
| Tris-HCl (pH 7.4) | 25 | B2005 (Tris)<br>J/4320/15 (HCl) | Melford (Tris)<br>Fisher (HCl) |
| NaF | 50 | S7920 | Merck |
| NaCl | 100 | S7653 | Merck |
| EDTA (pH 8) | 1 | E1644 | Merck |
| EGTA (pH 8) | 5 | E1102 | Melford |
| NaPPi | 10 | S6422 | Merck |
| Triton-X 100 | 1% | M143 | VWR |

#### Supplementary Table 4: Endogenous immunoprecipitation lysis buffer

| Compound: | Concentration (mM): | Catalogue number: | Supplier: |
|---|---|---|---|
| Sucrose | 320 | S0809 | Melford |
| EGTA | 1 | E0396-10G | Merck |
| Tris-HCl (pH 7.8) | 10 | B2005 (Tris)<br>J/4320/15 (HCl) | Melford (Tris)<br>Fisher (HCl) |
| β-merceptoethanol | 0.1% | M3148 | Merck |
| Sodium orthovanadate | 1 | S6508 | Merck |
| Benzamidine | 1 | 12072 | Merck |

| | | | |
|---|---|---|---|
| Phenylmethylsulphonyl fluoride | 0.1 | P7626 | Merck |

#### Supplementary Table 5: List of antibodies used for endogenous immunoprecipitation

| Target: | Host species: | Catalogue number: | Supplier: | Concentration: |
|---|---|---|---|---|
| TSPO | Goat | NB100-41398 | Novus | 2µg/mL (1:200) |
| CPT1a | Mouse | 66039-1-Ig | Proteintech | 4µg/mL (1:500) |
| - | Goat (IgG) | I-5000 | Vector | 2µg/mL |
| - | Mouse (IgG2A) | X0943 | Dako | 4µg/mL |

#### Supplementary Table 6: List of antibodies used for immunoblotting

| Target: | Host species: | Dilution: | Catalogue number: | Supplier: |
|---|---|---|---|---|
| TSPO | Rabbit | 1:1000, 2.5% (v/v) BSA in TBS-T (0.05%) | ab109497 | Abcam |
| CPT1a | Rabbit | 1:1000, 2.5% (v/v) BSA in TBS-T (0.05%) | 15184-1-AP | Proteintech |
| GLUT1 | Rabbit | 1:1000, 2.5% (v/v) BSA in TBS-T (0.05%) | 07-1401 | Merck |
| HK2 | Rabbit | 1:1000, 2.5% (v/v) BSA in TBS-T (0.05%) | 2867S | Cell Signalling Technologies |
| pNFκB (p65) (Ser536) | Rabbit | 1:1000, 2.5% (v/v) BSA in TBS-T (0.05%) | 3033S | Cell Signalling Technologies |
| VDAC1 | Rabbit | 1:1000, 2.5% (v/v) BSA in TBS-T (0.05%) | ab15895 | Abcam |
| GAPDH | Rabbit | 1:1000, 2.5% (v/v) BSA in TBS-T (0.05%) | G9545 | Merck |
| NFκB (p65) | Mouse | 1:1000, 2.5% (v/v) BSA in TBS-T (0.05%) | 6956S | Cell Signalling Technologies |
| Myc-tag | Mouse | 1:1000, 5% milk (v/v) in TBS-T (0.05%) | 66004-1-Ig | Proteintech |
| β-actin | Mouse | 1:10,000, 5% milk (v/v) in TBS-T (0.05%) | NB600-501 | Novus |

| | | | | |
|---|---|---|---|---|
| Mouse (AlexaFluor 680) | Goat | 1:10,000, 5% milk (v/v) in TBS-T (0.05%) | A21057 | Abcam |
| Rabbit (DyLight 800 Conjugated) | Goat | 1:10,000, 5% milk (v/v) in TBS-T (0.05%) | MX10 | Rocklands |

#### Supplementary Table 7: List of antibodies used for immunocytochemistry

| **Target:** | **Host species:** | **Dilution:** | **Catalogue number:** | **Supplier:** |
|---|---|---|---|---|
| GFAP | Mouse | 1:1000, 1% NDS in PBST | mab360 | Millipore |
| Mouse (AlexaFluor-488) | Donkey | 1:500, 1% NDS in PBST | A-21202 | Invitrogen |

### Supplementary Figures

#### Supplementary Figure 1: Genotyping TSPO$^{-/-}$-line MPAs

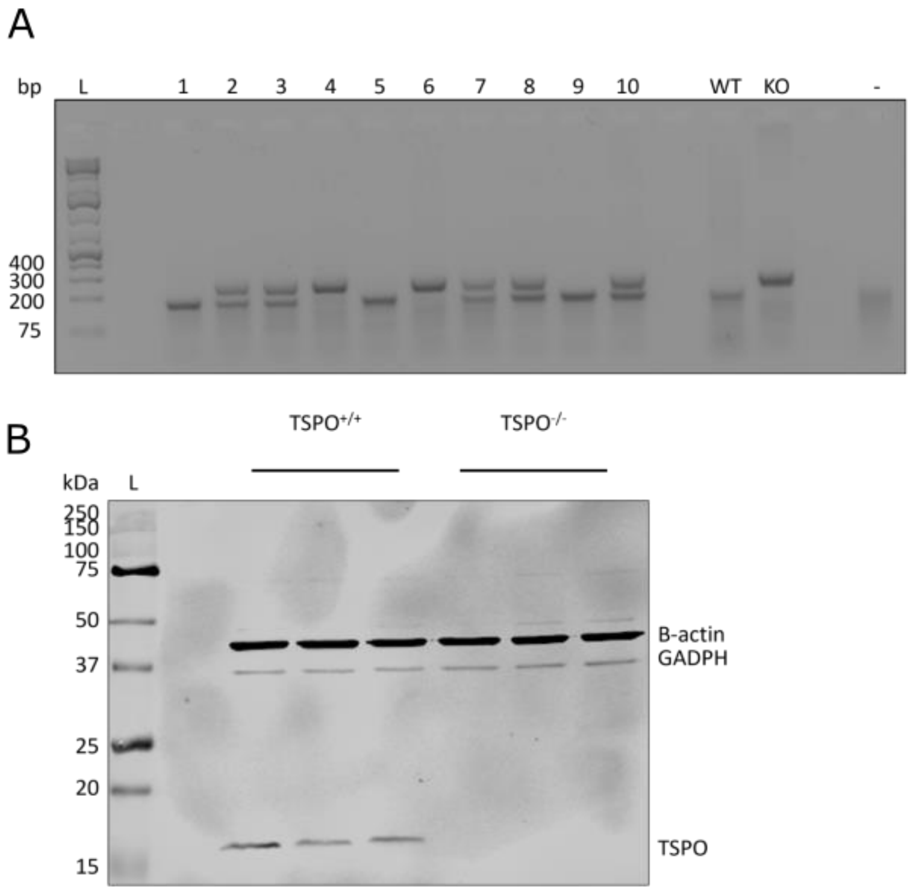


Genotypes of MPAs isolated from TSPO$^{-/-}$-line neonates were confirmed by PCR (**A**) and immunoblotting (**B**). **A:** Example image of a 1.5% agarose gel showing PCR products from TSPO$^{-/-}$-line neonates. TSPO$^{-/-}$ mice were indicated by the presence of a sole band at ~244kb. TSPO$^{+/+}$ mice were indicated by a sole band at ~188kb. TSPO$^{+/-}$ mice were not used for experiments, and were indicated by the presence of bands at ~244kb and ~188kb. **B:** Representative immunoblot of validation of functional TSPO knockout – TSPO (18kDa) was not detectable in lysate from TSPO$^{-/-}$ MPAs on immunoblots (abcam, ab109497).

#### Supplementary Figure 2: Confirmation of MPA purity

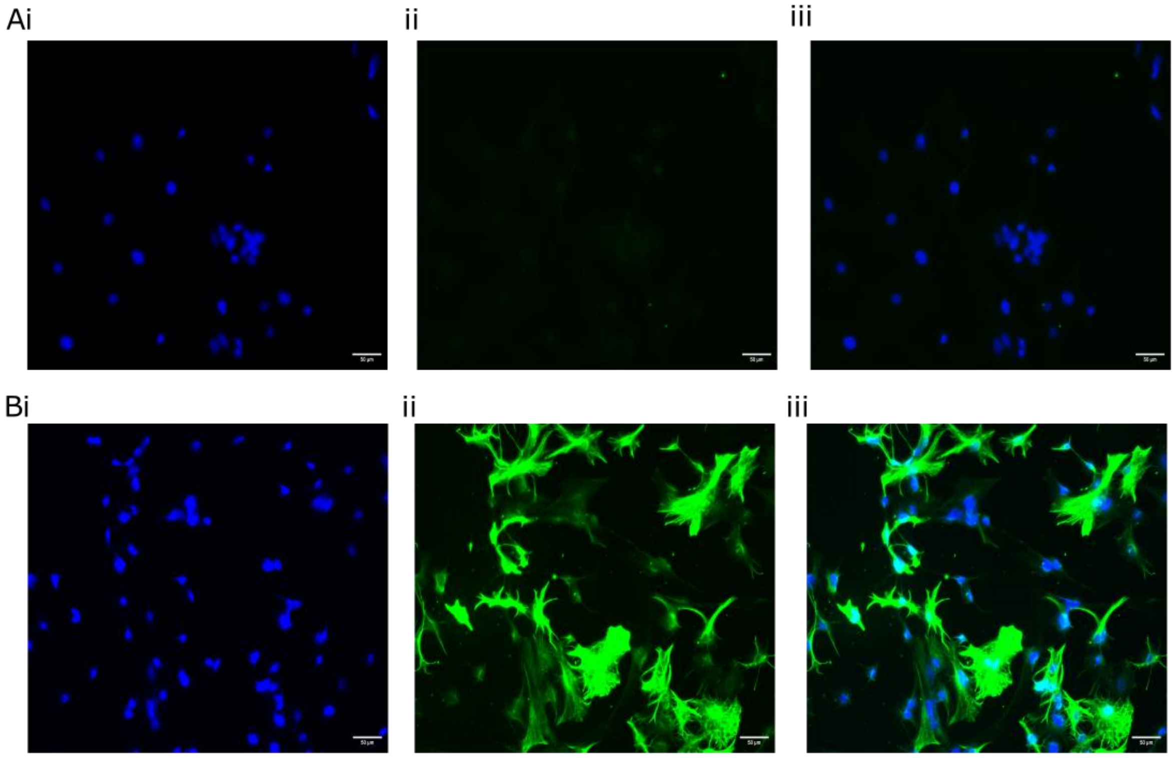


Purity of mouse primary astrocyte isolations was determined by immunocytochemistry. Cells were stained for glial fibrillary acidic protein (GFAP) and 4′,6-diamidino-2-phenylindole (DAPI, nuclear stain). A no-primary control was included in all stains to identify autofluorescence. Cells that were GFAP-positive were deemed to be astrocytes. Mean astrocyte purity was determined to be 98.67% (n=10 coverslips, 2-5 images per coverslip over 3-4 individual collections).

**A,Bi:** DAPI staining to identify nuclei. **Aii:** GFAP no-primary control. **Aiii:** Overlay of **Ai** and **Aii**. **Bii:** GFAP stain. **Biii:** Overlay of **Bi** and **Bii**.

Supplementary Figure 3: Confirmation of CRISPR-Cas9-generated TSPO genotype in U373 cells

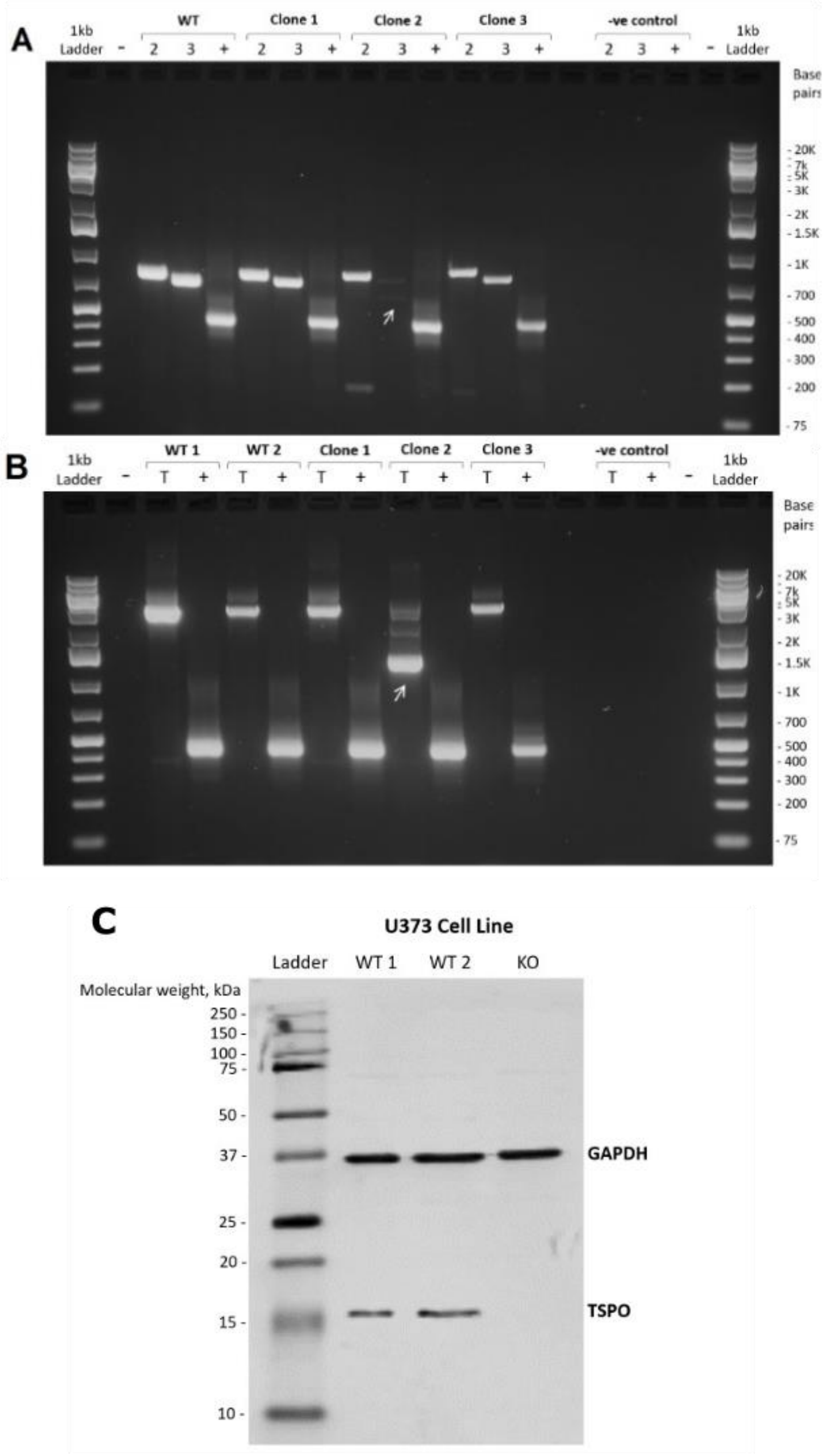

**A:** Confirmation from 3 CRISPR-Cas9 transfected U373 clones via PCR. Separate primer pairs were used to identify mutations in *Tspo* exons 2 and 3. **B:** PCR confirmation using a single pair of primers to amplify *Tspo* exons 2 and 3, confirming the mutation matches the predicted amplicon size of 1445bp, indicating a 1805bp deletion between target sites. **A,B:** Arrows indicate mutant amplicons of interest. WT, WT1: DNA from GM-naïve U373 cells. WT2: DNA from U373 cells transfected with empty plasmid. -ve control: DNA-free negative control. -: dd$H_2O$ loading control. 2: primer pair for *Tspo* exon 2. 3: primer pair for *Tspo* exon 3. T: primer pair for *Tspo* exons 2 and 3. +: positive control.

**C:** Representative immunoblot of confirmation of functional TSPO knockout in U373 cells. TSPO (abcam, ab109497) was not expressed in mutant cells, and was not altered by transfection with the empty vector plasmid. WT1: protein from GM-naïve U373 cells. WT2: protein from U373 cells transfected with the empty vector plasmid U373 cells. KO: protein from U373 knockout cells demonstrating TSPO deficiency.

Supplementary Figure 4: MPAs and U373 cells had different bioenergetic profiles

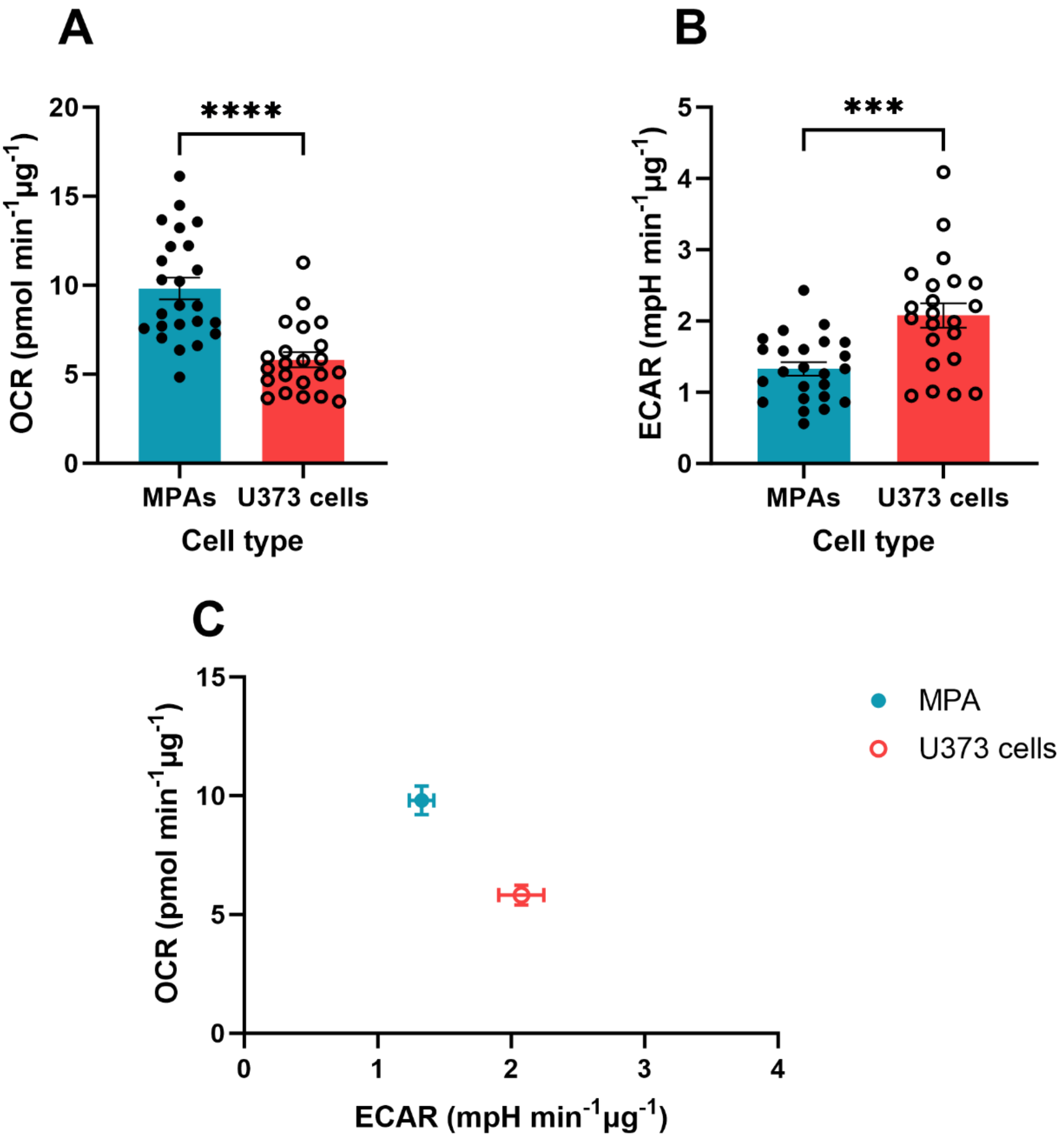


**A:** Oxygen consumption rate (OCR) of C57BL/6J MPAs compared to U373 cells. **B:** Extracellular acidification rate (ECAR) of C57BL/6J MPAs compared to U373 cells. **C:** Matrix showing OCR:ECAR comparison in MPAs and U373 cells.

**A:** Mann-Whitney test. **B:** Unpaired two-tailed t-test.

n=22-24. ***$p<0.001$, ****$p<0.0001$. Data are presented as mean ± standard error of the mean.

#### Supplementary Figure 5: 24h LPS stimulation did not significantly reduce MPA viability

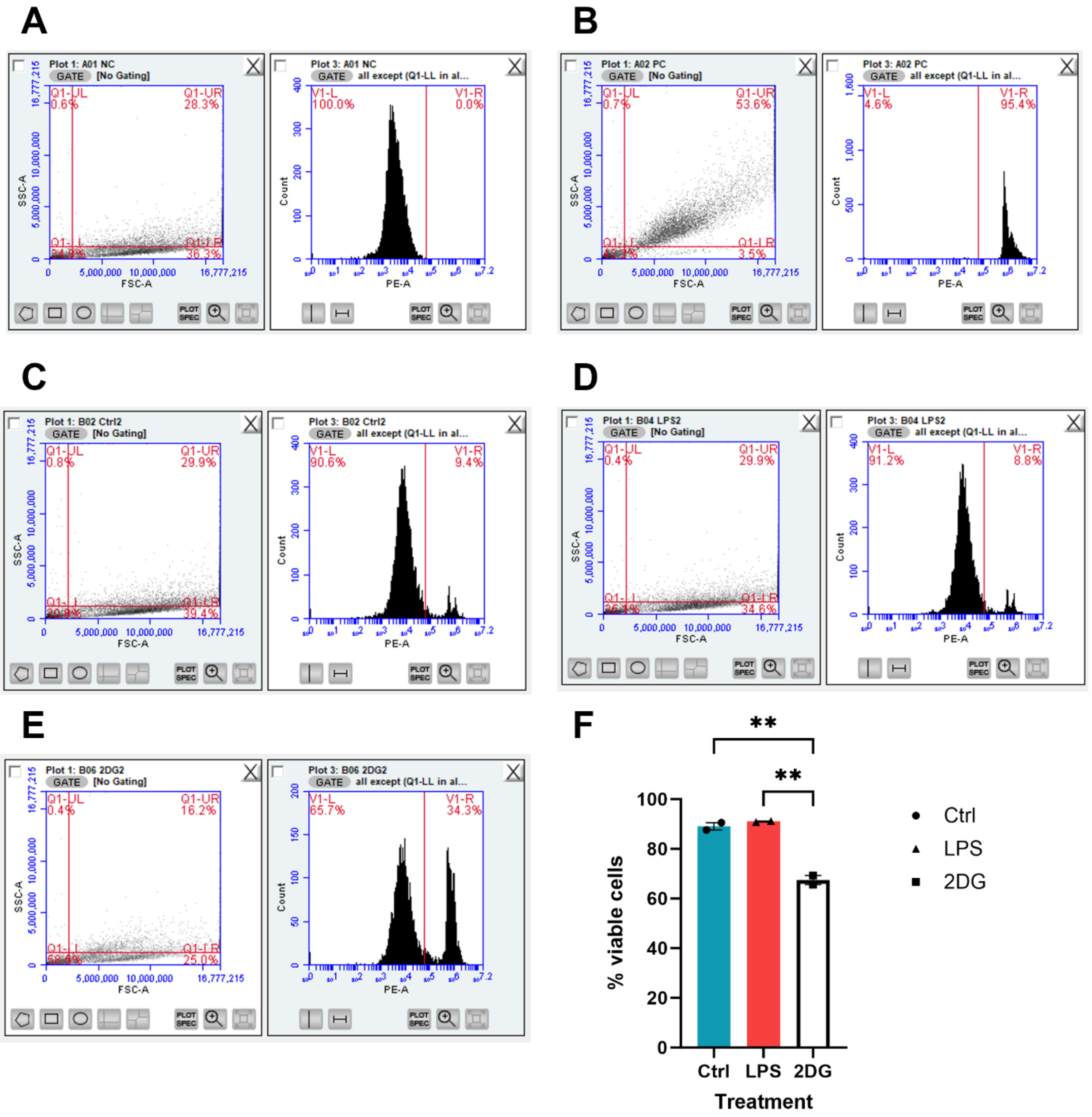


Settings used for gating flow cytometry for cell viability assay, measured by propidium iodide uptake, and example data. **A:** Negative control data used to gate experiment. Counts from Q1LL were deemed to be debris and excluded from counts. **B:** Example positive control data (MPAs boiled at 55°C for 10 minutes) to confirm validity of gating. **C:** Example data from control treated MPAs. **D:** Example data from 24h LPS stimulated MPAs. **E:** Example data from 24h 2DG stimulated MPAs. **F:** Cell viability data showing that 24h LPS stimulation did not significantly change MPA viability. 24h 2DG stimulation significantly reduced MPA viability.

n=2, one-way ANOVA with Tukey's multiple comparison's test. **p<0.01.

Ctrl: control. LPS: lipopolysaccharide, 100ng/mL. 2DG: 2-deoxyglucose, 10mM.
